# Rab32/Rab38-positive Lysosome-Related Organelle degrades lipid droplet in hepatocytes by microautophagy

**DOI:** 10.64898/2026.02.13.705687

**Authors:** Zidi Zhang, Shiou-Ling Lu, Yumiko Kato, Tongxin Zheng, Bohan Chen, Yangjie Li, Yu Usami, Taki Nishimura, Ryohei Sakai, Tomohiro Kabuta, Narikazu Uzawa, Satoru Toyosawa, Takeshi Noda

**Affiliations:** Department of Oral Cellular Biology, Center for Frontier Oral Science, Graduate School of Dentistry, The University of Osaka, Osaka, Japan; Department of Oral & Maxillofacial Oncology and Surgery, Graduate School of Dentistry, The University of Osaka, Osaka, Japan; Graduate School of Frontier Biosciences, The University of Osaka, Osaka, Japan; Center for Infectious Disease Education and Research, The University of Osaka, Osaka, Japan; Department of Oral and Maxillofacial Pathology, Osaka University Graduate School of Dentistry, Osaka, Japan; Institute for Protein Research, The University of Osaka, Osaka, Japan; Department of Degenerative Neurological Diseases, National Institute of Neuroscience, National Center of Neurology and Psychiatry, Tokyo Japan

**Keywords:** Rab32, Rab38, lysosome-related organelles (LROs), lipid droplet degradation, hepatic lipid metabolism, phosphatidylinositol 3-phosphate (PI3P), phosphatidylinositol 3, 5-bisphosphate (PI(35)P_2_), hepatocytes

## Abstract

Rab32 and Rab38 are paralogous small GTPases involved in the biogenesis of lysosome-related organelles (LROs), yet their roles in hepatic lipid metabolism remain poorly defined. Here, Rab32 and Rab38 double-knockout (DKO) male mice exhibited an age-dependent increase in body weight accompanied by hepatic lipid accumulation, suggesting impaired hepatic lipid processing. In AML12 hepatocytes, Rab32 and Rab38 localized to ring-like, LAMP1-positive structures characteristic of LROs, whose size increased with cell confluence.

Pharmacological inhibition of lysosomal acid lipase with orlistat led to the accumulation of lipid droplets (LDs) within Rab32/38-positive LROs, indicating that LD degradation occurs in these compartments. Additional treatment with bafilomycin A1 revealed invagination-like internal membrane structures within enlarged LROs. These processes were not affected by artificial inhibition of macroautophagy, highlighting the involvement of microautophagy. Ring-like signals positive for phosphatidylinositol 3-phosphate (PI3P) or phosphatidylinositol 3,5-bisphosphate (PI(3,5)P₂) were detected within or adjacent to LRO membranes, and LDs were frequently associated with these structures, suggesting a role for PI3P and PI(3,5)P₂ in internal membrane formation. Vps4B was also required for efficient LD incorporation. Consistently, Rab32/38 double-knockdown (DKD) AML12 cells exhibited increased lipid accumulation, indicating impaired LD engulfment.

Together, these findings identify Rab32/38-positive LROs as a structural platform for microautophagy-mediated lipid droplet degradation in hepatocytes.

## Introduction

Lipid droplets (LDs) are conserved intracellular organelles that store neutral lipids, primarily triacylglycerols and cholesteryl esters, and play essential roles in cellular energy balance, membrane synthesis, and lipid homeostasis (Henne et al., 2025; Olzmann & Carvalho, 2019; Walther & Farese, 2012). In hepatocytes, LD turnover is achieved through two major pathways: cytosolic lipolysis, mediated by enzymes such as adipose triglyceride lipase (ATGL) (Zechner et al., 2012), and lysosome-dependent degradation, which delivers LD-derived lipids into acidic compartments (Schulze et al., 2020; Singh et al., 2009). Although macroautophagy-dependent lipophagy was initially considered the major lysosomal route for LD degradation, emerging evidence indicates that LD processing involves multiple, mechanistically distinct lysosomal pathways. Recently, a LAMP2B-dependent direct LD-lysosome interaction pathway was identified in hepatocytes, revealing that LDs can be transferred to lysosomes through physical tethering independent of classical macroautophagy (Sakai et al., 2025). This study highlights the existence of specialized lysosomal subdomains dedicated to LD uptake and degradation, underscoring that LD clearance depends not only on canonical lysosomes but also on additional lysosome-related compartments whose molecular identity remains unclear.

Rab GTPases are master regulators of membrane trafficking, organelle identity, and cargo specificity (Stenmark, 2009). Among them, Rab32 and Rab38 are well recognized for their central roles in the biogenesis and maturation of melanosomes, a prototypical lysosome-related organelle (LRO) found in pigment cells (Bultema et al., 2014; Wasmeier et al., 2006). These GTPases direct cargo delivery, membrane remodeling, and acquisition of organelle-specific features, establishing melanosomes as functionally specialized LROs distinct from classical lysosomes.

Beyond pigment cells, Rab32/38-mediated LRO pathways have emerged as conserved mechanisms across diverse cell types. Previous studies demonstrated that Rab32 and Rab38 regulate intracellular membrane trafficking and LRO formation in osteoclasts and macrophages, contributing to specialized degradative or phagocytic functions (Noda et al., 2023; Tokuda et al., 2023). More recently, work from our laboratory extended the functional scope of Rab32/38-positive LROs in macrophages by demonstrating microautophagy-mediated mitochondrial degradation, in which LRO membranes invaginate to engulf damaged mitochondria. This finding establishes Rab32/38-associated LROs as active mediators of organelle quality control beyond classical lysosomal degradation pathways (Lu et al., 2025). These studies suggest that Rab32/38-dependent LRO biogenesis may operate in other tissues where specialized degradative compartments are required.

Despite these advances, the functions of Rab32 and Rab38 in hepatocytes remain largely unexplored. Although the liver relies heavily on lysosome-associated lipid degradation, it is unknown whether hepatocytes harbor Rab32/38-positive lysosome-related organelle (LRO)-like compartments and whether these organelles contribute to lipid droplet (LD) degradation. Furthermore, the mechanisms by which Rab32 and Rab38 regulate organelle identity, membrane organization, and interactions with LDs in hepatocytes have not been investigated.

In this study, we examine the involvement of Rab32 and Rab38 in LD degradation in hepatocytes. By defining the identity, dynamics, and lipid-processing functions of Rab32/38-positive LROs, we uncover a previously unrecognized pathway linking LRO biology to hepatic lipid metabolism.

## Materials and methods

### Cell Culture, Plasmids

Murine AML12 hepatocyte cells (ATCC, CRL-2254) were maintained in DMEM/F-12 supplemented with HEPES buffering system (11330032, Invitrogen; 500 mL), 10% fetal bovine serum (FBS; F7524, Sigma-Aldrich), 1× Penicillin-Streptomycin Solution (100×stock; 168-23191, Fujifilm Wako) and 1× Insulin-Transferrin-Selenium Solution (ITS-G) (100×stock; 41400045, Invitrogen) and 40 ng/ml Dexamethasone (≥98%, HPLC; D1756, Sigma-Aldrich) at 37 °C in a humidified 5% CO₂ incubator. For HaloTag labeling, cells were incubated in DMEM containing 100 nM TMR-conjugated Halo ligand (G8251, Promega) for 20 min, followed by washing with fresh medium before subsequent treatments. For experiments requiring defined cell confluence, AML12 cells were seeded at specified densities depending on the experimental setup. For biochemical analyses, including western blotting, cells were seeded in 6-well plates at densities of 1.25 × 10⁵ (low), 2.5 × 10⁵ (medium), or 5 × 10⁵ (high) cells per well and harvested the following day. For live-cell imaging experiments, cells were seeded on 35-mm glass-bottom dishes at densities of 1.5 × 10⁴ (low), 3.0 × 10⁴ (medium), or 6.0 × 10⁴ (high) cells per dish and cultured for 1-2 days prior to imaging. Incubation times were adjusted according to the experimental design and the reported growth characteristics of AML12 cells (Wu et al., 1994).

pLKO.1-puromycin plasmids expressing shRNAs were used for gene silencing. For *Rab32* and *Rab38* knockdown, shRNAs were designed in-house and cloned into the pLKO.1-puromycin vector. *Rab32* knockdown was achieved using an shRNA targeting nucleotide position 293 of the *Rab32* coding sequence (sh*Rab32*; target sequence: 5′-GGACTTCGCCCTCAAAGTTCT-3′). *Rab38* knockdown was achieved using an shRNA targeting nucleotide position 321 of the *Rab38* coding sequence (sh*Rab38*; target sequence: 5′-GGACATTGCTGGTCAAGAAAG-3′). For *Vps4a* and *Vps4b* knockdown, pLKO.1-puromycin plasmids expressing shRNAs targeting mouse *Vps4a* and *Vps4b* (TRCN0000101416 and TRCN0000101824) were purchased from Sigma-Aldrich. The pLKO.1-puro control plasmid was a gift from Professor Makoto Sato (Graduate School of Medicine, The University of Osaka). pMRX-puro-*Lamp1*-mRFP, pMRX-puro-EGFP-*Rab32* and -*Rab38* plasmids were prepared as described previously (Etoh & Fukuda, 2019). Mouse *Rab5A* and *Rab7* cDNAs were subcloned into the pMRX-bsr-mStr plasmid or pMRX-puro-mStr vector (Etoh & Fukuda, 2019; Noda et al., 2023) from the constructs pMRX-puro-EGFP-*Rab5A* and pMRX-puro-EGFP-*Rab7* (Tsuboi & Fukuda, 2006).

The mCherry-EGFP-PLIN2 (Sakai et al., 2025) and PX-SnxA^GV^-GCC-mCherry (Nishimura et al., 2025) were previously described. Halo-LC3B (#207539), mCherry-2×FYVE (#140050) and mStrawberry-Atg4B^C74A^ (#21076) were obtained from Addgene. All constructs were sequence validated. Plasmid transfections were performed using Lipofectamine™ 2000 Transfection Reagent (11668019, Thermo Fisher Scientific) following the manufacturer’s instructions.

### Antibodies

Primary and secondary antibodies used in this study were as follows. LC3 was detected using rabbit anti-LC3 (PM036, MBL; 1:500 for IF and 1:2000 for WB). p62/SQSTM1 was detected with rabbit anti-p62 (PM045, MBL; 1:2000 for WB). Rab GTPases were detected using mouse anti-Rab32 (sc-390178, Santa Cruz Biotechnology; 1:100 for IF and 1:200 for WB) and mouse anti-Rab38 (sc-390176, Santa Cruz Biotechnology; 1:100 for IF and 1:200 for WB). Tubulin was detected using mouse anti-tubulin (66031-1-Ig, Proteintech; 1:10,000 for WB). LAMP1 was detected using rat anti-LAMP1 (1D4B) (sc-19992, Santa Cruz Biotechnology; 1:100 for IF). S6 kinase was detected using rabbit anti-phospho-S6K (Thr389) (#9205S, Cell Signaling Technology; 1:1000 for WB) and rabbit anti-S6K (#9202, Cell Signaling Technology; 1:1000 for WB). Secondary antibodies for immunofluorescence included Alexa Fluor 488- and 568-conjugated anti-rabbit IgG (A11008 and A10042, Thermo Fisher Scientific; 1:1000), Alexa Fluor 488 and 568-conjugated anti-mouse IgG (A11001 and A10037, Thermo Fisher Scientific; 1:1000), and Alexa Fluor 488-conjugated anti-rat IgG (A21208, Thermo Fisher Scientific; 1:1000). For immunoblotting, HRP-conjugated anti-rabbit IgG (111-035-003, Jackson ImmunoResearch; 1:5000) and anti-mouse IgG (115-035-003, Jackson ImmunoResearch; 1:5000) were used.

### Chemical Reagents

Oleic acid (O1257, Sigma-Aldrich; stock: 9 mM in sterile water; working concentration: 200 µM) was used to induce lipid droplet formation. Orlistat (S1629, Selleck Chemicals; stock: 100 mM in DMSO; working concentration: 50 or 200 μM) was used to block lysosomal acid lipase activity; bafilomycin A1 (BVT-0252-M001, BioViotica; stock: 500 µM in DMSO; working concentration: 20 nM, 100 nM or 200 nM) was used to inhibit the acidification of lysosome-related organelles (LROs); Apilimod (STA5326, Selleck Chemicals; stock: 10 mM in DMSO; working concentration: 100 nM) was used to inhibit PIKfyve activity; SAR405 (HY-12481, MedChemExpress; stock: 10 mM in DMSO; working concentration: 1 μM) added immediately after transient transfection and maintained until completion of live-cell imaging to suppress Vps34 activity. Lipid droplet dyes included Lipi-Blue and Lipi Deep Red (LD01 and LD04, Dojindo Laboratories; stock: 0.1 mM; working concentration: 0.1 µM), while the acidified areas in LROs were dyed with LysoTracker Red DND-99 (L7528, Thermo Fisher Scientific, stock: 100 µM in DMSO; working concentration: 75 nM). Puromycin (P8833, Sigma-Aldrich; stock: 2 mg/mL in sterile water; working concentration: 2 μg/ml) and blasticidin S (KK-400, Kanto Chemical Co., Inc.; stock: 10 mg/mL in sterile water; working concentration: 5∼10 μg/ml) were used for the selection of stable cell lines.

### Mouse model

Wild-type C57BL/6J mice were purchased from Japan SLC (Shizuoka, Japan). *Rab32/38* double knockout C57BL/6J mice were described previously (Tokuda et al., 2023). Mice were housed in a specific pathogen-free (SPF) facility under controlled environmental conditions (22–24 °C, 40–60% humidity) with a 12 h light/12 h dark cycle, and were provided ad libitum access to sterilized chow diet and autoclaved water in the animal facility of the Graduate School of Dentistry, The University of Osaka. Unless otherwise specified, all experiments were performed using male mice. Both male and female mice were used for body weight analyses. All procedures were approved by the Institutional Animal Experiments Committee of The University of Osaka Graduate School of Dentistry (29-008-0, R-06-003-0) and the Gene Modification Experiments Safety Committee of The University of Osaka.

### Retroviral Production and Overexpression Constructs

Retroviral particles for overexpression of GFP-Rab32, GFP-Rab38, LAMP1-RFP, mStr-Rab5A, mStr-Rab7 or mRFP-LC3 were generated in Platinum-E (Plat-E) packaging cells (Morita et al., 2000). Plat-E cells were transfected with pMRX-puro-EGFP-Rab32, pMRX-puro-EGFP-Rab38, pMRX-puro-LAMP1-RFP, pMRX-bsr-mStr-Rab5A, pMRX-puro-mStr-Rab7, or pMRX-bls-mRFP-LC3 using PEI (Polyethyleneimine) (Boussif et al., 1995), following previously described retroviral production procedures. Viral supernatants were collected at 48 h and 72 h post-transfection, passed through 0.45-µm PVDF syringe filters (Millex-HV, SLHVR33RS, Merck Millipore), supplemented with 8 μg/mL polybrene (hexadimethrine bromide; H9268, Sigma-Aldrich; stock: 10 mg/mL in sterile water), and used to infect AML12 cells. Infected cells were selected with puromycin (2 µg/mL) for 5-7 days to establish stable overexpression lines. For the generation of dual-expressing AML12 cell lines (e.g., GFP-Rab32/LAMP1-RFP and GFP-Rab38/LAMP1-RFP), the fluorescence double positive cells were isolated by fluorescence-activated cell sorting (FACS) using a Sony SH800S cell sorter.

### Lentiviral Production and Stable shRNA Knockdown

Lentiviral particles were generated in HEK293 cells using PEI as the transfection reagent. For the generation of lentiviral particles, Halo-LC3B or pLKO.1-puro vectors encoding sh*Rab32*, sh*Rab38*, sh*Vps4a*, or sh*Vps4b* were co-transfected with the packaging plasmids psPAX2 and pMDG at a 1:3 DNA-to-PEI ratio in serum-free medium. After incubation for 15-20 min at room temperature, the DNA-PEI complexes were added dropwise to HEK293 cells. Virus-containing supernatants were collected and used for cell transduction following the same procedure as retrovirus system. Knockdown efficiency was confirmed by immunoblotting or qPCR.

### qPCR

Cell pellets were treated with TRIsure (BIO-38032, Bioline) to extract total mRNA, and the iScript cDNA Synthesis Kit (1725038, Bio-Rad) to generate cDNA. Real-time PCR analysis was performed with the Step One Plus Real-time PCR System (Applied Biosystems) using THUNDERBIRD^®^ SYBR^®^ qPCR Mix (QPS-201, TOYOBO) and the primers were designed based on reference mRNA sequences from the NCBI RefSeq database, for VPS4A (NM_126165.2): ACGGTGGAATGATGTAGCTGG (forward), CCAAAGAGGAGTATGCCTCGC (reverse); VPS4B (NM_009190.2): GGCTGCACGGAGAATTAAGAC (forward), TCCAGAACCCAGGGTATATTTGT (reverse); GAPDH (NM_008084.4): AAATGGTGAAGGTCGGTGTG (forward), TGAAGGGGTCGTTGATGG (reverse). Relative gene expression levels for each gene were calculated according to the ΔΔCt method. The knockdown efficiency of shRNA-targeted genes was normalized to the gene levels in the scramble control.

### Western blotting

AML12 cells were cultured in 6-well plates at defined seeding densities (1.25 × 10⁵, 2.5 × 10⁵, and 5 × 10⁵ cells per well) to achieve low, medium, and high confluence, respectively. For experiments requiring larger protein amounts, cells were cultured in 6-cm dishes. At the indicated time points, cells were placed on ice, washed three times with ice-cold PBS (137 mM NaCl, 2.7 mM KCl, 10 mM Na₂HPO₄, 1.76 mM KH₂PO₄, pH 7.4), and detached using a sterile 13-mm cell scraper (90020, SPL Life Sciences). Suspensions were transferred to 1.5-mL tubes and pelleted at 900 × g for 3 min at 4 °C. Pellets were resuspended in ice-cold lysis buffer [50 mM Tris-HCl (pH 7.5), 150 mM NaCl, 1 mM DTT, 1% Triton X-100, protease inhibitor cocktail (11873580001, Roche)] and incubated on ice for 20 min. Lysates were cleared by centrifugation at 20,400 × g for 10 min at 4 °C. Supernatants were mixed with 6× SDS sample buffer (300 mM Tris-HCl pH 6.8, 12% SDS, 30% glycerol, 0.006% bromophenol blue, 0.6 M 2-mercaptoethanol) and boiled for 5 min at 95 °C. Equal protein amounts were resolved by SDS-PAGE (10-15% gels) and transferred onto PVDF membranes (IPVH00010, Millipore). Membranes were blocked in 5% skim milk in PBS-T (0.05% Tween20 in PBS; Tween20: polyoxyethylene (20) sorbitan monolaurate; 166-21115, FUJIFILM Wako) and incubated overnight at 4 °C with primary antibodies as described above. After washing, membranes were incubated with HRP-conjugated anti-mouse or anti-rabbit IgG. Signals were developed using Immobilon® Forte Western HRP Substrate (Millipore, WBLUF0100), captured on a GeneGnome-5 imaging system (Syngene Bio Imaging), and quantified using ImageJ (NIH).

### Immunofluorescence Staining and Confocal Microscopy

Cells were cultured on 12-mm glass coverslips (No. 1S thickness; C012001, Matsunami Glass) placed in 24-well plates. Cells were fixed by adding 4% paraformaldehyde (PFA, 163-20145, Fujifilm Wako) for 5 min, replaced with fresh 4% PFA for an additional 5 min at room temperature (RT), and then washed with PBS. Permeabilization was performed using 50 μg/mL digitonin (300410, Calbiochem) in PBS for 10 min at RT, followed by blocking with 0.2% gelatin (076-02765, Fujifilm Wako) in PBS for 30 min.

Cells were incubated with primary antibodies against target proteins (e.g., Rab32, Rab38, LAMP1, LC3B; see Antibody part) diluted in blocking buffer, followed by Alexa Fluor-conjugated secondary antibodies (Thermo Fisher Scientific) for 1 h at RT. After final washes, coverslips were mounted using ProLong Glass Antifade Mountant (P36984, Thermo Fisher Scientific). For fluorescent probe-based imaging, dyes such as Lipi-Blue and Lipi Deep Red (LD01 and LD04, Dojindo Laboratories), LysoTracker Red DND-99 (L7528, Thermo Fisher Scientific), or dextran-Rhodamine B, 70,000 MW, neutral (D1841, Molecular Probes) were applied according to the manufacturer’s instructions prior to fixation or during live imaging as appropriate. For live-cell imaging, cells were cultured on 35-mm glass-bottom dishes (D11350H, Matsunami Glass) and imaged at 37 °C using the heated stage of the Leica TCS SP8 confocal microscope. Confocal images were acquired using a Leica TCS SP8 laser scanning microscope equipped with an HC PL APO CS2 63×/1.40 NA oil-immersion objective (Leica Microsystems). Imaging parameters, including laser power, detector gain, and pinhole size, were kept constant across experimental conditions. Fluorescence intensity, vesicle size, and vesicle number were quantified using ImageJ.

### Dextran Long-Pulse Endocytic Labeling

To visualize fluid-phase uptake and label lysosome-related organelles (LROs), AML12 cells were incubated with 0.25 mg/mL dextran-Rhodamine B, 70,000 MW, neutral (D1841, Molecular Probes) for 18-24 h at 37 °C. After labeling, cells were washed three times with warm medium and immediately imaged by confocal microscopy. Long-pulse labeling ensured dextran accumulation in late endocytic and LRO compartments.

### High-Fat Diet Feeding and Tissue Collection

Wild-type and Rab32/38 DKO mice were fed a high-fat diet (45% kcal fat; D12451, Research Diets) beginning at 5 weeks of age and continued for 12 weeks. Body weight was measured every week. Mice were euthanized after the indicated feeding period, and liver and adipose tissues were harvested, weighed, and processed for biochemical analysis. Blood samples were collected by cardiac puncture following overnight fasting for serum triacylglycerol measurement.

### Triacylglycerol Measurement

Serum triacylglycerol levels of mice fed a normal chow diet were measured by an external laboratory service (Oriental Yeast Co., Ltd., Osaka, Japan). For mice fed a high-fat diet (HFD), serum triacylglycerols were quantified according to the manufacturer’s instructions using the LabAssay™ Triglyceride kit (291-94501, FUJIFILM Wako).

Liver triacylglycerol content was determined following a modified Folch lipid extraction method (Folch et al., 1957). In brief, approximately 20-30 mg of liver tissue was homogenized in sodium chloride buffer and extracted with chloroform/methanol (2:1, vol/vol). After phase separation, the organic layer was collected and dried under nitrogen gas, and triacylglycerol concentrations were determined using the same assay. Values were normalized to tissue weight.

### Histology

Tissues were fixed overnight in 4% PFA at 4 °C, embedded in paraffin, and sectioned at 5 µm thickness. Sections were stained with hematoxylin and eosin (H&E) for morphological assessment. Berlin Blue staining for the detection of iron deposition in liver sections was performed using the Berlin Blue Staining Set (296-21541, FUJIFILM Wako) following the protocol described previously (Nikiforov, 2017). Stained sections were imaged using a Nikon Eclipse Ci microscope equipped with a 40 × objective lens.

### Statistical Analysis

Statistical analyses were performed using GraphPad Prism 9.5.1 for macOS. Data are presented as mean ± SD or mean ± SE, as indicated in the corresponding figure legends. For comparisons between two groups, Student’s t-test was used when data were normally distributed with equal variances, and Welch’s t-test was applied when variances were unequal. For datasets that did not follow a normal distribution, the Mann-Whitney U test or Kolmogorov-Smirnov test was used. For Western blot quantification normalized to a reference condition (e.g., low-confluence samples normalized to 1), one-sample t-tests were performed to compare medium- or high-confluence values against the theoretical value of 1. For comparisons among more than two groups, one-way ANOVA was applied when assumptions of normality and homoscedasticity were satisfied; when these assumptions were violated, Welch’s ANOVA or appropriate non-parametric alternatives were used. For experiments involving two factors, two-way ANOVA or mixed-effects models (REML) were applied depending on the variance structure and the presence of missing values. Post hoc tests (Tukey’s, Dunnett’s, or Šidák’s) were used when appropriate with adjustments for multiple comparisons. Exact statistical tests and n values are provided in the figure legends. \*\*\*\**p* < 0.0001; \*\*\**p* < 0.001; 0.001 < \*\**p* < 0.01; and 0.01 < \**p* < 0.05 in figures show the level of significance.

## Results

### Expression level of Rab32 and Rab38 affects the size of LRO in AML12 cells

AML12 cell, a mouse hepatocyte line widely used for studying lipid metabolism (Das et al., 2024; Sakai et al., 2025; Schott et al., 2019; Schulze et al., 2020), was immuno-stained with Rab32 or Rab38 together with LAMP1, and their colocalization on ring-like organelle indicates their identity as lysosome-related organelle (LRO) (Fig. 1A, B) (Noda et al., 2023; Tokuda et al., 2023). Larger ring-like LAMP1-positive LROs were co-localized with GFP-Rab32/38, whereas smaller LAMP1-positive punctuated structures did not, suggesting that GFP-Rab32/38-positive LROs are distinctive from conventional LAMP1-positive lysosomes as in macrophage and osteoclast (Fig. S1A) (Noda et al., 2023; Tokuda et al., 2023). Quantitative analysis showed that the colocalization of GFP-Rab32/38 with LAMP1-RFP remained unchanged across experimental conditions, including those described in later sections (Fig. S1B). Rab32/38-positive LROs extensively overlapped with Rab7, a late endosome marker, whereas only partially with Rab5, an early endosome marker (Fig. 1C-E), suggesting that Rab32/38-positive LROs share features with late endosomes. Co-localization of endogenous Rab32 with GFP-Rab38, and of endogenous Rab38 with GFP-Rab32, suggests their overlapping function as reported in macrophage, osteoclast, and melanocyte (Fig. 1F) (Noda et al., 2023; Tokuda et al., 2023; Wasmeier et al., 2006).

**Figure 1.**
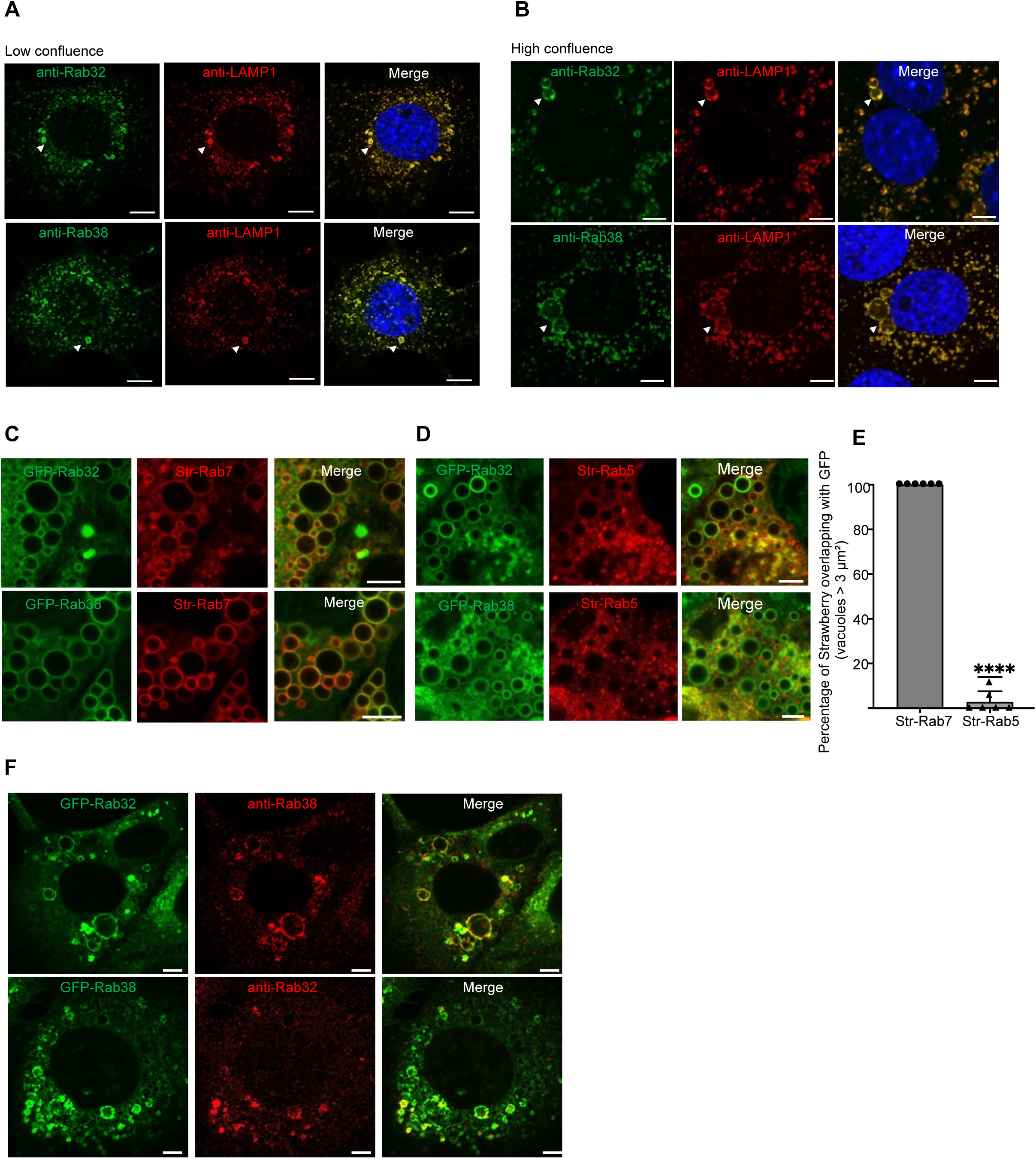
Characteristics of Rab32/38-positive LROs. **A, B**. Representative immunofluorescence images of AML12 cells at low (30%) and high (80%) confluence stained for Rab32 or Rab38 (green) and LAMP1 (red). Nuclei were stained with DAPI (blue). Arrowheads indicate colocalization of Rab32/38 with LAMP1. Scale bars, 5 μm (low confluence) and 7.5 μm (high confluence). **C.** Representative live-cell images of GFP-Rab32 or GFP-Rab38 with Strawberry-Rab7 in AML12 cells. Scale bar, 5 μm. **D.** Representative live-cell images of GFP-Rab32 or GFP-Rab38 with Strawberry-Rab5 in AML12 cells. Scale bar, 5 μm. **E.** Quantification of colocalization between GFP-Rab38-positive LROs (>3 μm²) and Strawberry-Rab7 or Strawberry-Rab5 (n = 5 random fields per condition). Statistical significance was determined using a one-sample t-test (*p* < 0.0001). Scale bar, 5 μm. **F.** Representative immunofluorescence images showing GFP-Rab32 with endogenous Rab38 and GFP-Rab38 with endogenous Rab32. Scale bar, 5 μm.

LRO was visualized with Rhodamine B-conjugated dextran (MW ∼70 kDa) which was added to the medium and incorporated via fluid-phase endocytosis. We noticed that the number of large Rhodamine-positive vacuoles, whose volume is more than 3 µm^2^, increased when cell density reached high confluence (∼80%) compared to the condition in low confluence (∼30%) (Fig. 2A, B). At low confluence, smaller Rab32/38-positive LROs were predominant, whereas at high confluence larger LROs were frequently observed (Fi g. 1A, B). Both Rab32 and Rab38 are expressed in AML12 cells, and it is of note that relative expression levels of each protein gradually increased as cell confluence progressed from ∼30% to ∼80% (Fig. 2C, D). To ask their role in LRO formation, Rab32 and Rab38 were knocked down with shRNA in AML12 cells (Fig. 2E). Rab32 expression showed a modest increase upon Rab38 knockdown, suggesting their compensation (Fig. 2E). Compared with control AML12 cells at high confluence, double knockdown (DKD) cells displayed a marked reduction in large LRO number (Fig. 2F, G). Knockdown of Rab38 alone tended to reduce vacuole numbers like that observed in DKD cells, whereas Rab32 knockdown had less impact on vacuole formation (Fig. S1C). Conversely, overexpression of GFP-Rab38 at low confluence increased large vacuoles and GFP-Rab32 exerted a similar but less efficient effect than Rab38 (Fig. 2H, I). Collectively, these results indicate that the protein level of Rab38 and Rab32 plays a crucial role in determining the size of LRO.

**Figure 2.**
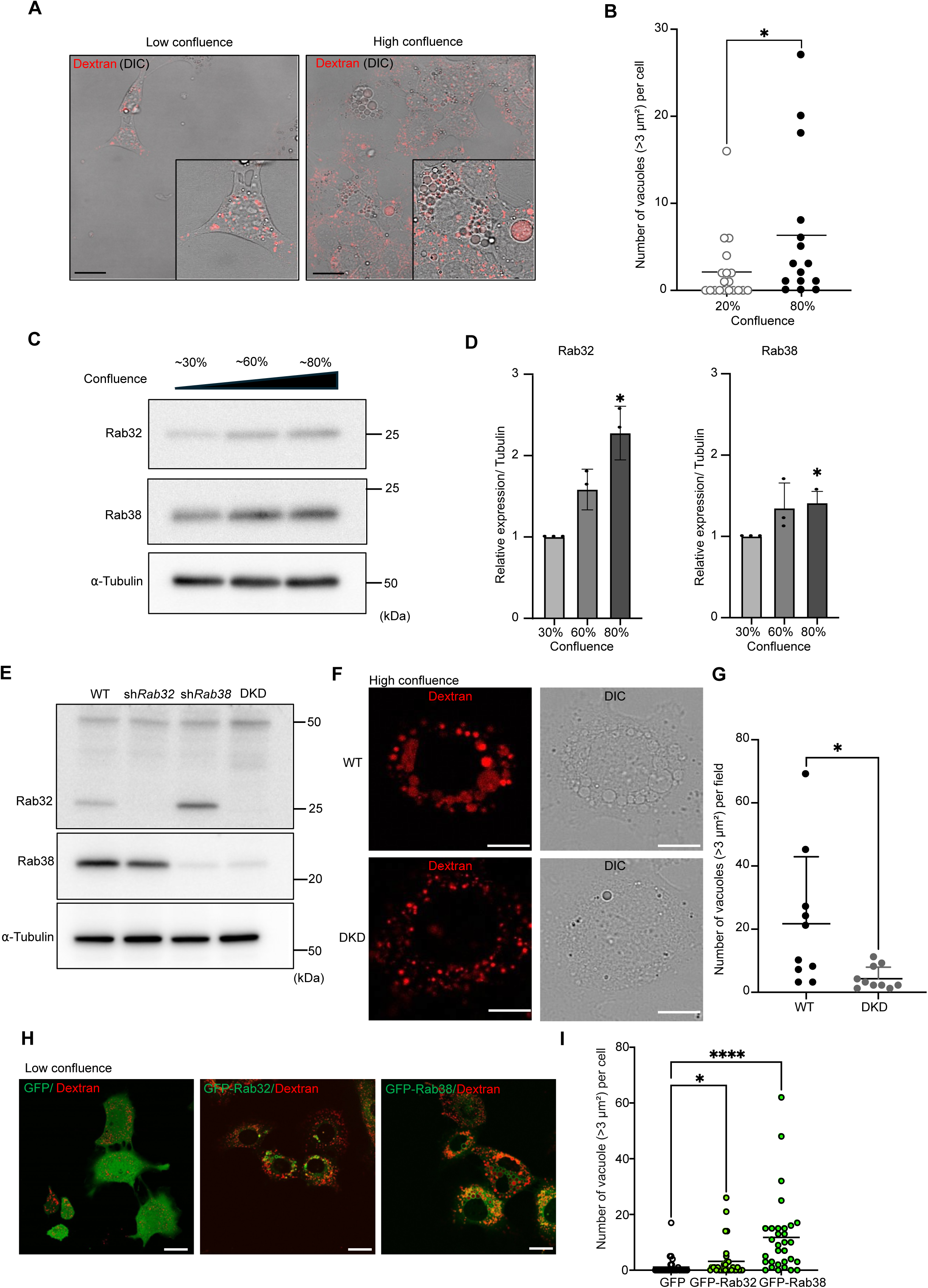
Rab32 and Rab38 are required for large LRO formation in AML12 cells. **A.** Representative DIC and dextran staining images of AML12 cells at low (30%) and high (80%) confluence. Scale bar, 25 μm. **B.** Quantification of vacuole (> 3 μm²) number per cell. Data are mean ± SD (low, n = 19; high, n = 15). Statistical significance was determined using the Mann-Whitney test (p = 0.0415). **C.** Immunoblotting of Rab32 and Rab38 in AML12 cells at ∼30%, ∼60%, and ∼80% confluence. Tubulin was used as a loading control. **D.** Quantification of Rab32 and Rab38 protein levels relative to tubulin. Bars represent mean ± SD from three independent experiments. *p* values were determined by one-sample *t*-test; Rab32: 30% vs 60% (*p* = 0.0561), 30% vs 80% (*p* = 0.0215). Rab38: 30% vs 60% (*p* = 0.1924), 30% vs 80% (*p* = 0.0389). **E.** Immunoblot analysis of Rab32 and Rab38 expression in control, shRab32, shRab38, and Rab32/38 double knockdown (DKD) AML12 cells. Tubulin was used as a loading control. A nonspecific band at ∼50 kDa was detected with the anti-Rab32 antibody. **F.** Representative DIC and dextran staining images of WT and DKD AML12 cells at high confluence. Scale bar, 10 μm. **G.** Quantification of vacuole number (>3 μm²) per field in WT and DKD cells (n = 10 random fields per condition, 5 cells per field). Data are mean ± SD. Statistical significance was determined using the Mann-Whitney test (*p* = 0.0071). **H.** Representative live-cell imaging of dextran-stained vacuoles in AML12 cells expressing GFP control, GFP-Rab32, or GFP-Rab38 at low confluence. Scale bar, 25 μm. **I.** Quantification of vacuole number (>3 μm²) per single cell (n = 25 cells per condition). Data are mean ± SD. Statistical significance was determined using the Mann-Whitney test (GFP vs GFP-Rab32, *p* = 0.0304; GFP vs GFP-Rab38, *p* < 0.0001).

### Rab32/38-positive LROs mediate lipid droplet degradation independently of macroautophagy

Next, whether Rab32/38-positive LROs are involved in lipid droplet (LD) dynamics was investigated. AML12 cells expressing both GFP-Rab32 and LAMP1-RFP were used for live-cell imaging and cultured under standard growth conditions and stained with Lipi-Blue, the dye specifically stains lipid droplets (Tatenaka et al., 2019), and LDs were rarely observed inside LRO (Fig. 3A, upper panel). However, if cells were treated with orlistat, the lipase inhibitor (Tuohetahuntila et al., 2017), LD could be frequently observed inside the LRO, suggesting that these LRO constitutively incorporate LD to be degraded (Fig. 3A, middle panel). Notably, quantitative analysis showed that orlistat treatment did not affect the colocalization of GFP-Rab32/38 with LAMP1-RFP, as comparable levels of colocalization were observed in the presence or absence of orlistat (Fig. S1B). To examine the involvement of LRO membrane remodeling in this incorporation process, bafilomycin A1, an inhibitor of V-ATPase that suppresses acidification of lysosomes and LROs, thereby inhibiting intraluminal disintegration (Noda et al., 2023), was employed. Combined treatment with bafilomycin A1 and orlistat revealed that LDs were surrounded by LAMP1-positive membranes through either simple membrane invaginations or more complex multilayered structures (Fig. 3A, bottom panel). These observations suggest that LD incorporation within LROs involves extensive membrane remodeling. Oleic acid is supplied in the culture medium to enhance LD formation (Fujimoto & Parton, 2011; Listenberger et al., 2003). Oleic acid supplementation not only increased LD abundance but also markedly enhanced the formation of invagination-like membrane structures surrounding LDs within GFP-Rab32-positive LROs (Fig. S1D), further supporting the notion that LD incorporation into LROs is accompanied by active membrane remodeling.

**Figure 3.**
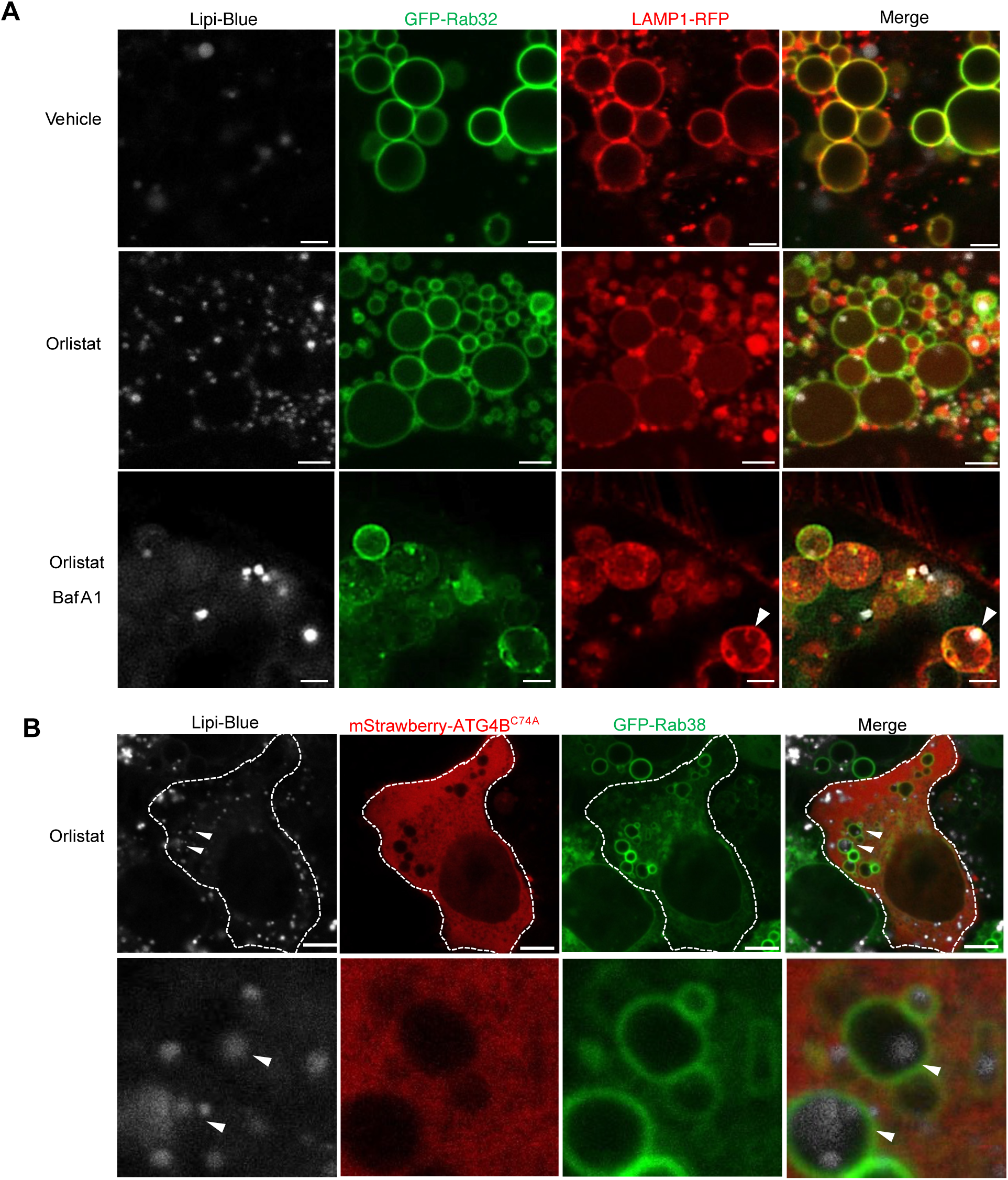
Rab32/38-positive LRO-mediated lipid droplet degradation is independent of macroautophagy. **A.** Representative live-cell fluorescence images of AML12 cells expressing GFP-Rab32 and LAMP1-RFP, stained with Lipi-Blue (white), under vehicle, 50 µM orlistat, or 50 µM orlistat and 20 nM Bafilomycin A1 (BafA1) for 24 h. Arrowheads indicate lipid droplets enclosed by Rab32/38-positive LROs. Scale bar, 2.5 µm. **B.** Representative images of AML12 cells treated with 50 µM orlistat and co-expressing mStrawberry-Atg4BC74A and GFP-Rab38. Lipi-blue marks lipid droplets. Dashed lines outline cell boundaries. Arrowheads indicate LDs engulfed by Rab38-positive LROs, indicating macroautophagy-independent LD engulfment. Scale bar, 5 µm.

There were several reports that the macroautophagic process plays a role in delivering LD to lysosome to be degraded (Schulze et al., 2020; Singh et al., 2009; Zechner et al., 2012). AML12 cells displayed high autophagic flux under nutrient-rich conditions (Fig. S2A), and consistent with previous reports, a fraction of LDs localized adjacent to LC3-positive puncta (Wang et al., 2017); however, a substantial subset of LDs was closely associated with GFP-Rab32/38 positive LROs without detectable LC3 signals in their vicinity (Fig. S2B). To ask whether this incorporation process depends on macroautophagy, a previously established system, that macroautophagy is completely suppressed, was employed (Fujita et al., 2008). ATG4B is a processing proteinase for LC3 family members essential for macroautophagy, and overexpression of its inactive mutant results in inhibition of LC3 lipidation, thereby blocking macroautophagy (Fig. S2C) (Fujita et al., 2008). As expected, in ATG4B inactive mutant (C74A)-overexpressing cells, LC3 puncta, representing autophagosome formation, failed to form under starvation confirming effective inhibition of macroautophagy (Fig. S2D). Remarkably, despite this macroautophagy-deficient condition, LDs were still observed within LROs following orlistat treatment (Fig. 3B). These results indicate that LD degradation in Rab32/38-positive LROs occurs independent of macroautophagy.

### Rab32 and Rab38 differentially regulate LD morphology and lysosomal delivery

To determine how Rab32 and Rab38 influence lipid droplet (LD) dynamics, we quantified total LD area per cell in Rab32 or Rab38 single-knockdown (SKD) and double-knockdown (DKD) AML12 cells. Knockdown of Rab32 or Rab38 individually, as well as double knockdown of both, led to lipid droplet accumulation. (Fig. 4A, B). Interestingly, the morphology of accumulated LDs differed between the knockdowns: *Rab32* SKD cells preferentially accumulated enlarged LDs, whereas *Rab38* SKD cells accumulated numerous small LDs (Fig. S3A-D).

**Figure 4.**
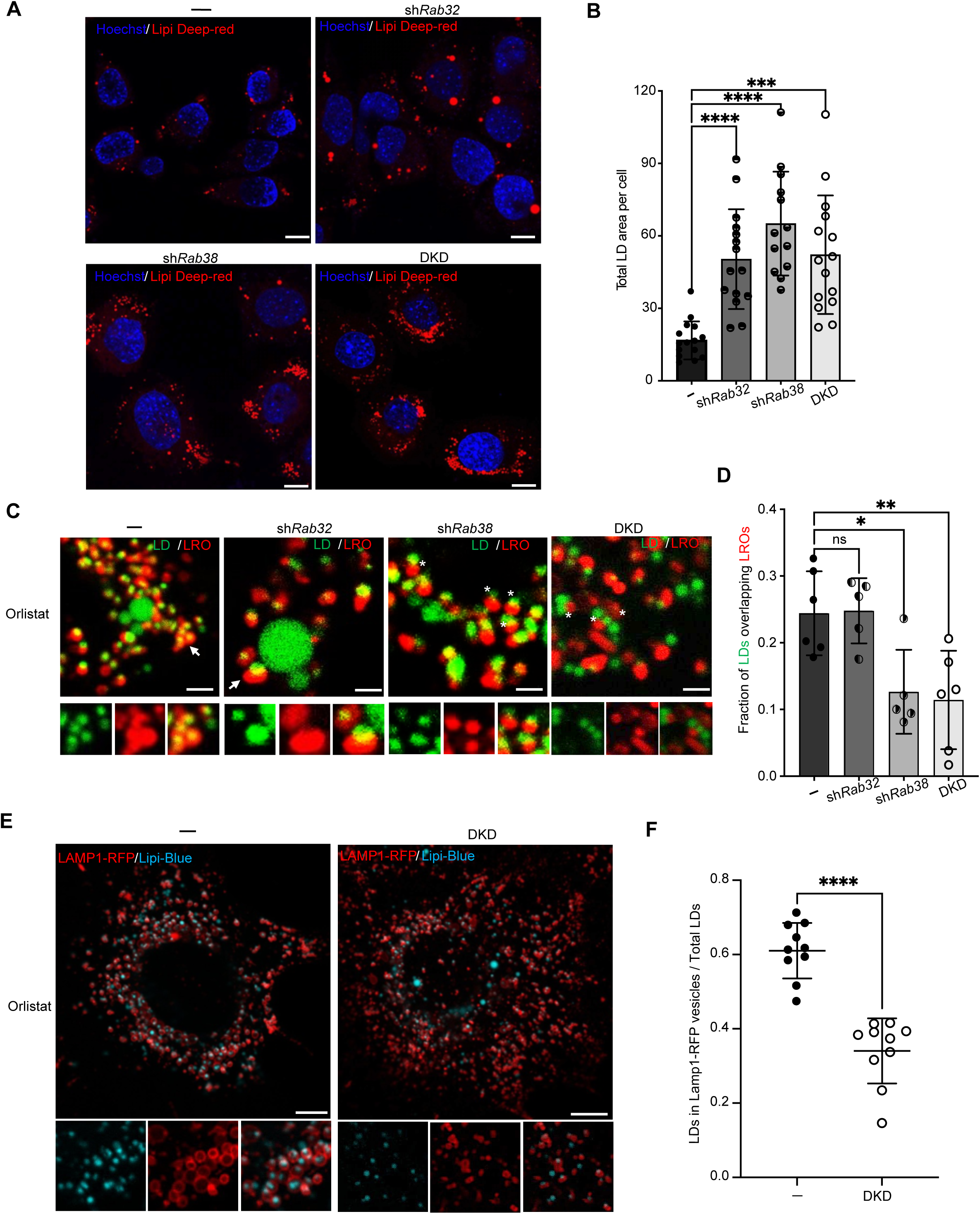
Loss of Rab32/38 leads to impaired lipid droplet engulfment by LROs. **A.** Representative fluorescence images of AML12 control (-), sh*Rab32*, sh*Rab38*, and *Rab32/38* double-knockdown (DKD) cells stained with Hoechst (blue) and Lipi Deep-Red (red). Scale bars, 10 µm. **B.** Quantification of total LD area per cell in control (-), sh*Rab32*, sh*Rab38*, and DKD cells. Data are presented as mean ± SD (control, n = 15; sh*Rab32*, n = 15; sh*Rab38*, n = 13; DKD, n = 15). Statistical significance was determined using Brown-Forsythe ANOVA with multiple comparisons (control vs sh*Rab32*, *p* < 0.0001; control vs sh*Rab38*, *p* < 0.0001; control vs DKD, *p* = 0.0003). **C.** Representative images of orlistat-treated control (-), sh*Rab32*, sh*Rab38*, and DKD cells stained with Lipi-Blue (green) and Lysotracker (red). Arrowheads indicate LDs surrounded by lysosomes, whereas asterisks indicate LDs showing impaired lysosomal engagement in DKD cells. Insets show magnified views. Scale bar, 5 µm. **D.** Quantification of the fraction of LDs overlapping with lysosomal structures in each genotype. Data are mean ± SD from randomly acquired fields (control, n = 6; sh*Rab32*, n = 5; sh*Rab38*, n = 5; DKD, n = 6). Statistical significance was determined using ordinary one-way ANOVA with multiple comparisons (control vs sh*Rab32*, ns, *p* = 0.993; control vs shRab38, *p* = 0.0181; control vs DKD, *p* = 0.0064). **E.** Representative images of WT and DKD AML12 cells expressing LAMP1-RFP (red) and stained with Lipi-blue (cyan) following orlistat treatment. Insets show zoomed regions highlighting LD-lysosome associations. Scale bar, 10 µm. **F.** Quantification of the proportion of LDs localized inside LAMP1-RFP positive structures relative to total LDs in WT and DKD cells. Data are mean ± SD from 10 cells in each group. Statistical significance was determined using the Mann-Whitney test (*p* < 0.0001).

To further evaluate LD-LRO interactions, cells were treated with orlistat. In control and *Rab32* SKD cells, LDs were frequently detected inside LysotrackerRed-positive LROs. In contrast, LDs were rarely observed inside acidified compartments in *Rab38* SKD and DKD cells (Fig. 4C, D). To validate this finding with an orthogonal system, we employed a mCherry-GFP-mPLIN2 reporter targeted to LDs (Fig. S4A), in which GFP fluorescence is quenched upon entry into acidic environments, but not mCherry signal, allowing selective visualization of LDs delivered to acidic organelles (Sakai et al., 2025; Schulze et al., 2020). As a result, DKD cells showed significantly fewer mCherry-only signals, supporting reduced lysosomal delivery of LD (Fig. S4B, C). It is noteworthy that Lysotracker signal intensity was diminished in DKD cells, raising the possibility that acidification of LRO is impaired, as observed in macrophages and osteoclasts (Noda et al., 2023), and could affect pH-dependent probes. Therefore, orthogonal analysis was performed by generating a LAMP1-RFP overexpression cell line. As result, fewer LDs were engulfed by LAMP1-positive structures in DKD cells compared with control AML12 cells (Fig. 4E, F). Consistent with independence from macroautophagy, *Rab32/38* double-knockdown cells exhibited no obvious impairment in LC3 flux, as assessed by Halo-LC3 (Yim et al., 2022) (Fig. S4D).

Together, these findings indicate that Rab32 and Rab38 both contribute to LD delivery into LROs.

### PI3P and PI(3,5)P₂ support membrane remodeling within Rab32/38-positive LROs

Phosphoinositides are major determinants of the identity of membrane organelle including those in the endosomal-lysosomal system (Posor et al., 2022), and PI3P is enriched on early and late endosomes, whereas PI(3,5)P₂ localizes to later-stage compartments such as multivesicular bodies and lysosomes (Volpicelli-Daley & De Camilli, 2007). PI3P dynamics during LD incorporation were visualized using an mCherry-tagged 2×FYVE probe. FYVE-positive signals frequently appeared as ring-like structures either within LROs or closely associated with their limiting membranes (Fig. 5A). Upon orlistat treatment, LDs were often observed inside or adjacent to FYVE-positive structures, which were themselves enclosed by the LRO membrane (Fig. 5A). When the cells were treated with SAR405, a highly selective inhibitor of Vps34/Class III PI3 kinase (Ronan et al., 2014), FYVE-positive rings almost completely disappeared (Fig. 5A). This treatment excluded the entry of LDs into LROs and resulted in accumulation outside of them (Fig. 5A), indicating that PI3P is required for LD enclosure by LRO membranes.

**Figure 5.**
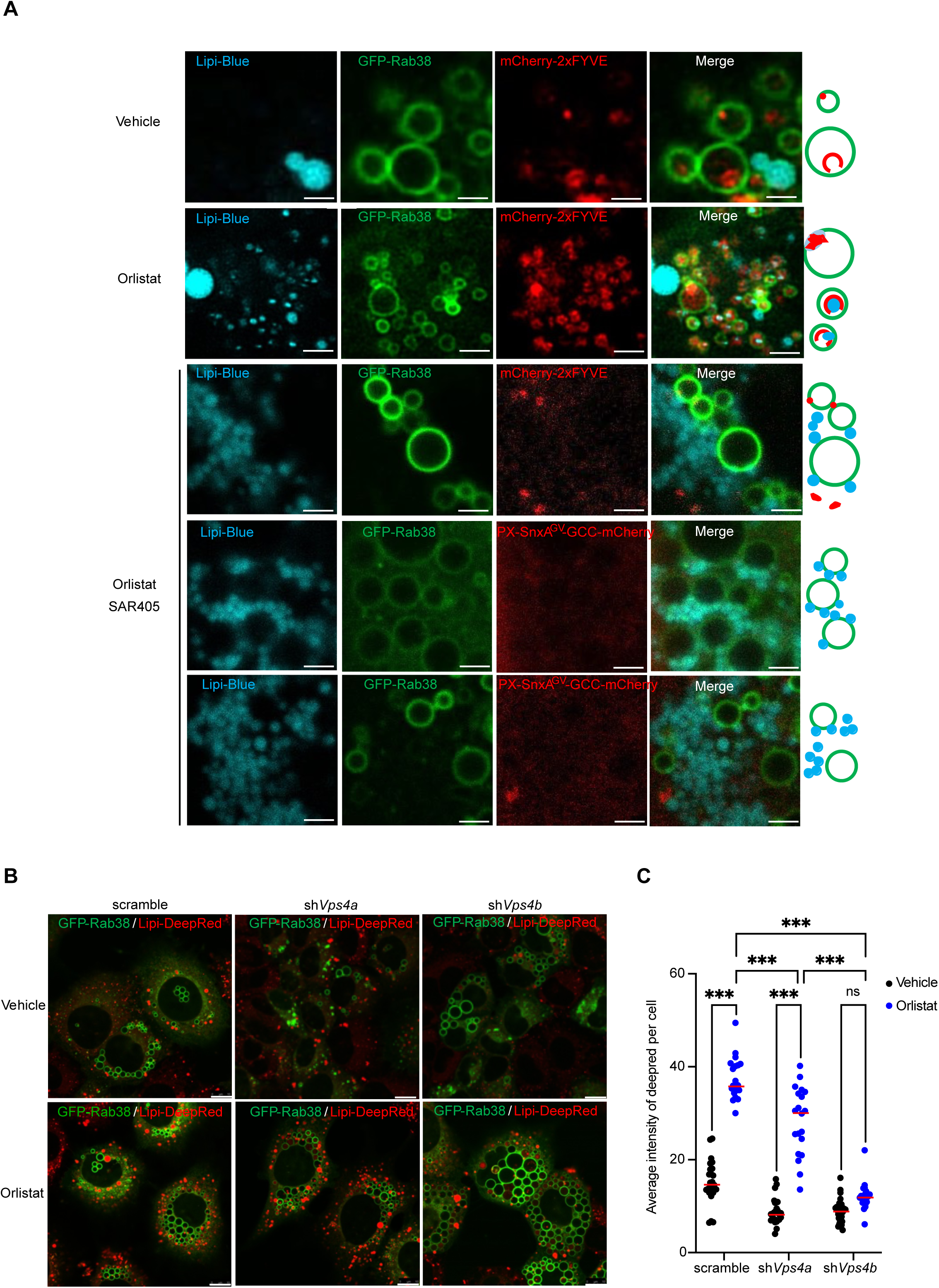
PI3P and PI(3,5)P₂ support membrane remodeling within Rab32/38-positive LROs, and VPS4B is required for lipid droplet delivery. **A.** Representative live-cell fluorescence images of AML12 cells expressing GFP-Rab38 together with the PI3P probe mCherry-2xFYVE or the PI(3,5)P₂ probe PX-SnxA^GV^-GCC-mCherry, stained with Lipi-Blue. Cells were treated with vehicle, 50 µM orlistat, or 50 µM orlistat and 1 µM SAR405 for 24 h. Scale bar, 2 µm. **B.** Representative live-cell fluorescence images of AML12 scramble, sh*Vps4a*, and sh*Vps4b* cells transduced with a GFP-Rab38-expressing retrovirus and stained with Lipi-DeepRed. Cells were treated with vehicle (DMSO) or orlistat (200 µM, 24 h) prior to imaging. Scale bar, 10 µm. **C.** Quantification of Lipi-DeepRed average intensity in scramble, sh*Vps4a*, and sh*Vps4b* AML12 cells transduced with GFP-Rab38 and treated with vehicle (DMSO) or orlistat. Data are presented as mean ± SD (n ≥ 20 single cells per condition). Statistical significance was determined by two-way ANOVA with multiple-comparisons testing. **** *p* < 0.0001; ns, not significant.

To further assess phosphoinositide involvement, a recently developed highly sensitive PI(3,5)P₂ probe was used (PX-SnxA^GV^-GCC-mCherry) (Nishimura et al., 2025). To acutely inhibit PI(3,5)P₂ synthesis, cells were treated with apilimod, a selective inhibitor of the PIKfyve, which phosphorylates PI3P to PI(3,5)P₂ (Cai et al., 2013; Jefferies et al., 2008). Under basal conditions, in the absence of apilimod treatment, small PX-SnxA^GV^-GCC-mCherry positive ring-like structures were occasionally observed adjacent to the LRO membrane, whereas in other LROs the probe signal appeared as diffuse or smeared mCherry fluorescence within the lumen (Fig. S5A). Following apilimod treatment, cells became dominated by markedly enlarged LROs, and the probe positive ring structures were no longer detectable (Fig. S5A). Upon apilimod washout, newly generated signals could be detected (Fig. S5B), and subsequent orlistat treatment induced the formation of the probe-positive ring-like assemblies that attached to lipid droplets and anchored them to the LRO membrane (Fig. S5B). This phenomenon was abolished by SAR405 treatment, suggesting that both PI3P and downstream PI(3,5)P₂ contribute to LRO membrane dynamics required for LD uptake (Fig. 5A). Together, these results indicate that PI3P and PI(3,5)P₂-dependent membrane remodeling generates internal membrane structures within Rab32/38-positive LROs, enabling efficient LD capture and degradation.

### VPS4B is required for LD delivery to LRO

ESCRT is a conserved membrane-remodeling machinery, including VPA4A/B, that mediates membrane scission events required for multivesicular body biogenesis, cytokinetic abscission, and plasma membrane repair (Vietri et al., 2020). Knockdown of *Vps4a/b* reduces lipid hydrolysis in AML12 cells, as well as other ESCRT components such as TSG101, ALIX, or CHMP4B, suggesting an ESCRT-dependent pathway for lipid degradation (Sakai et al., 2025). Moreover, previous work has demonstrated that VPS4A depletion selectively impairs LD degradation within LAMP2-positive lysosomes, whereas loss of VPS4B blocks LD degradation at Rab7-positive endosomes (Das et al., 2024).

Knockdown of *Vps4b* resulted in the formation of enlarged GFP-Rab38 positive LROs compared with scramble control cells (Fig. 5B). Whole-cell average lipid droplet (LD) fluorescence intensity was reduced in sh*Vps4b* cells (Fig. 5B, C), raising the possibility that cytosolic lipase activity may compensate for VPS4B loss. Upon orlistat treatment, LDs accumulated within GFP-Rab38 positive LROs in scramble cells, whereas LDs were rarely detected within Rab38-positive compartments in sh*Vps4b* cells (Fig. 5B, C), indicating a reduced association between LDs and GFP-Rab38 positive LROs under *Vps4b*-deficient conditions. In contrast, knockdown of *Vps4a* produced a comparatively milder phenotype. Taken together, these results suggest that VPS4B is essential for coupling lipid droplet cargo to Rab32/38-positive LROs.

### Rab32/38 deficiency causes age-dependent fat accumulation and increases susceptibility to diet-induced obesity

*Rab32/38* double-knockout (DKO) mice were generated previously (Tokuda et al., 2023). The male mice showed increased body weight at 12 months, but that is not the case of female mice (Fig. 6A). Consistent with this, 1-year-old DKO males displayed increased abdominal white adipose tissue (WAT) accumulation (Fig. 6B). Rab32 was expressed in liver and WAT but was nearly undetectable in brown adipose tissue (BAT) (Fig. 6C). Rab38 was detectable in WAT, but barely in liver and BAT (Fig. 6C). No significant weight change was observed in *Rab32* SKO or *Rab38* SKO mice of either sex (Fig. 6A, and S6A), indicating that the phenotype is dependent on the combined loss of Rab32 and Rab38.

**Figure 6.**
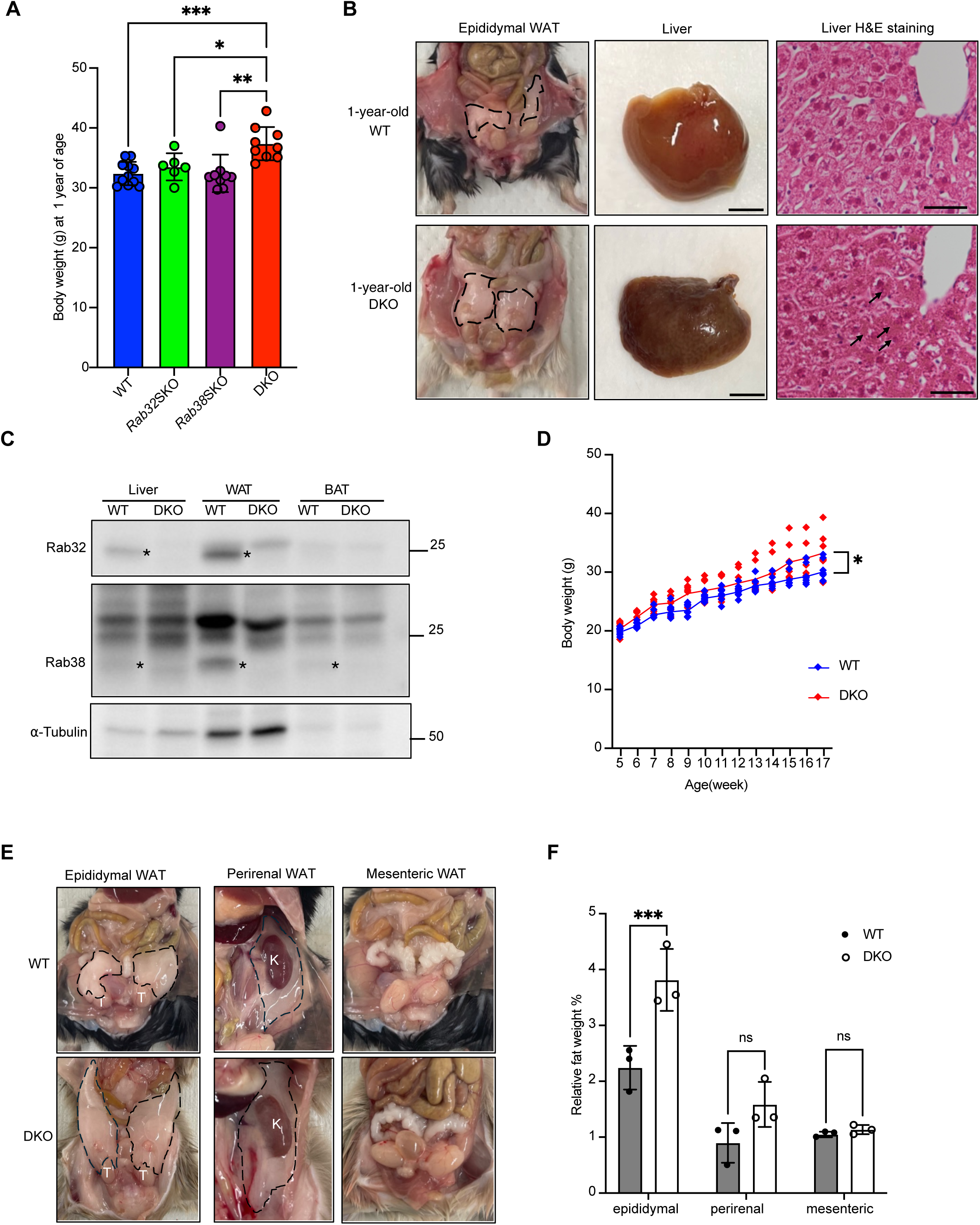
Rab32/38 deficiency causes age-dependent fat accumulation and increases susceptibility to diet-induced obesity. **A.** Body weight at 1 year of age of male WT, Rab32/38 double-knockout (DKO), Rab32 single-knockout (SKO), and Rab38 single-knockout (SKO) mice under chow feeding. Data are presented as mean ± SD (n ≥ 6 mice per genotype). Each data point represents the body weight of an individual mouse. Statistical significance was determined by one-way ANOVA with multiple-comparisons testing. \*\*\**p* < 0.001; \*\**p* < 0.05 and \**p* < 0.01. **B.** Representative images of epididymal white adipose tissue (WAT), gross morphology, and H&E staining of livers from 1-year-old WT and DKO mice. Dashed lines indicate the boundaries of adipose tissue. (WT, n = 3; DKO, n = 3). Scale bars: gross images, 5 mm; histology, 25 µm. **C.** Immunoblot analysis of Rab32 and Rab38 in liver, white adipose tissue (WAT), and brown adipose tissue (BAT) from WT and DKO mice. α-Tubulin was used as a loading control for each tissue. Asterisks indicate the specific bands corresponding to Rab32 and Rab38 at the expected molecular weights. **D.** Body-weight trajectories of WT and DKO male mice fed a 45% kcal high-fat diet. Each data point represents the body weight of an individual mouse at the indicated age, and lines indicate group means (WT, n = 8 mice; DKO, n = 8 mice). Statistical significance was determined using two-way ANOVA (*p* = 0.0191). **E.** Representative images of epididymal, perirenal, and mesenteric WAT in WT and DKO mice. Dashed lines indicate adipose depots. T, testis; K, kidney. **F.** Quantification of epididymal, perirenal, and mesenteric WAT weights normalized to body weight (WT, n = 3; DKO, n = 3). Statistical analysis was performed using two-way ANOVA test; epididymal WAT, *p* < 0.001; perirenal WAT, *p* = 0.0812 (ns); mesenteric WAT, ns (*p* > 0.05). Data are presented as mean ± SD.

The livers of male DKO mice appeared markedly darker than wild-type controls and Hematoxylin and Eosin staining showed brown cytoplasmic granules (Fig. 6B), which were consistent with lipofuscin, an undegradable aggregate of oxidized lipids and proteins that accumulates when lysosomal clearance is impaired, reflecting reduced lysosomal degradation capacity (Brunk & Terman, 2002). These granules were negative for Berlin Blue staining that stain hemosiderin, iron granules, indicating that the darker liver coloration was not due to iron deposition (Fig. S6A).

Because the phenotype emerged relatively late in aged mice under a normal chow diet, next the male mice were fed with a high-fat diet (HFD) to accelerate lipid accumulation. Following HFD feeding, DKO males gained more weight than WT controls (Fig. 6D). Abdominal WAT accumulation was greatly amplified under HFD (Fig. 6E, F), and hepatic TAG levels were further elevated in DKO mice (Fig. S6B, C). Together, these data indicate that Rab32 and Rab38 are required for normal lipid regulation, and their loss leads to age-dependent fat accumulation and increased sensitivity to dietary stress.

## Discussion

This study identifies Rab32/38-positive lysosome-related organelles (LROs) as key organelles mediating lipid droplet (LD) degradation mediated by microautophagy in AML12 cells. A defining advance of this study is the introduction of lysosome-related organelles (LROs) into the field of hepatocyte lipid metabolism. Previous studies have proposed lysosomal and vacuolar compartments as destinations for lipid droplets in hepatocytes (Schulze et al., 2020; Singh et al., 2009). However, lysosomes are generally defined as much smaller than prominent vacuolar structures observed in AML12 cells, with their appearance becoming more pronounced as cell confluence increased (Huotari & Helenius, 2011; Luzio et al., 2007; Saftig & Klumperman, 2009). We demonstrate that these large vacuolar structures are positive for Rab32/38, establishing them as LROs, guided by our previous work in osteoclasts and macrophages (Lu et al., 2025; Noda et al., 2023; Tokuda et al., 2023). This finding provides a conceptual link between hepatocytes and LRO biology and opens new avenues for understanding organelle specialization in hepatic lipid metabolism. This study reveals that LRO formation depends on the expression levels of Rab32 and Rab38 (Fig. 2C, D-I).

Building on the previous study, which relied primarily on pharmacological inhibitors of macroautophagy (Sakai et al., 2025), we further strengthened the conclusion that LD degradation in AML12 cells occurs independently of macroautophagy by employing a robust macroautophagy-inhibition system that we had previously established (Fig. 3B). Furthermore, culturing cells in fatty acid-free medium enabled clearer discrimination of individual lipid droplets and facilitated direct observation of lipid droplet incorporation into invaginating LROs via microautophagy (Fig. S1D).

These LROs engulfed LDs through a PI3P- and PI(3,5)P₂-dependent process, as demonstrated by the inhibitory effects of specific pharmacological inhibitors (Fig. 5A). Notably, LD-incorporating subdomains of Rab32/38-positive LRO membranes were selectively enriched in PI3P, even though these LROs were largely devoid of Rab5 (Fig. 1C–E), suggesting that PI3P is locally generated at sites of LD incorporation. In addition, PI(3,5)P₂ was enriched at these subdomains (Fig. S5B), indicating that PI3P is locally converted to PI(3,5)P₂ during the microautophagic process. Such spatially restricted phosphoinositide remodeling resembles that observed during microautophagy-mediated mitochondrial engulfment into LROs in macrophages (Nishimura et al., under revision). One possible mechanism is that PIKfyve is recruited to these sites through its FYVE domain–mediated binding to PI3P; this possibility warrants further investigation.

This study showed VPS4B plays a prominent role in LD entry into Rab38-positive LROs, whereas VPS4A appears to contribute more modestly (Fig. 5B, C). The enlargement of GFP-Rab38 positive LROs observed upon VPS4B depletion suggests that VPS4B-dependent ESCRT activity is required to coordinate lipid droplet incorporation with membrane remodeling events at LROs. This phenotype is consistent with a role for VPS4B in ESCRT-mediated microautophagy-like processes operating at Rab7-positive endosomal (Das et al., 2024) or LRO membranes (this study). By contrast, the comparatively milder phenotype associated with *Vps4a* knockdown indicates that VPS4A is not the primary ESCRT ATPase driving LD entry into Rab38-positive LROs in this context. Instead, VPS4A appears to function in a mechanistically distinct and context-dependent manner. Recent work has identified VPS4A as a selective receptor for LC3-dependent lipophagy in hepatocytes (Das et al., 2024). Together with our observations, these findings support a model in which VPS4A and VPS4B regulate LD turnover through parallel but non-redundant pathways, operating at different membrane compartments and utilizing distinct molecular mechanisms.

Interestingly, despite the tendency of hepatic lipid accumulation, serum TAG levels were no different in DKO mice compared to WT mice after HFD (Fig. S6B, C). One possible explanation is that impaired lipid droplet degradation in hepatocytes limits efficient mobilization and export of TAG into the circulation, thereby lowering serum TAG levels while promoting hepatic lipid retention.

Although AML12 cells exhibit prominent LROs, it is noteworthy that healthy hepatocytes in the liver *in vivo* do not display comparably large vacuolar structures (Goldblatt & Gunning, 1984; Łysek-Gładysińska et al., 2024; Wang & Boyer, 2004). Furthermore, Rab32 is readily detectable in mouse liver tissue, whereas Rab38 expression is minimal or undetectable compared with AML12 hepatocytes (Fig. S6D), suggesting that baseline Rab38 availability may partly determine susceptibility to vacuolation. Thus, AML12 cells should be regarded as a hepatocyte-derived cell line that exhibits modified and specialized characteristics relative to hepatocytes *in vivo*.

Previous studies have suggested that hepatocyte vacuolation can be influenced by intracellular glycogen content, which alters the appearance and dynamics of vacuolated structures under metabolic stress (Nayak et al., 1996). This raises the possibility that metabolic state contributes to the extent of vacuolation observed in AML12 cells. However, the activity of mTORC1, a master regulator of cellular metabolic status (Laplante & Sabatini, 2012; Saxton & Sabatini, 2017; Wullschleger et al., 2006), remained comparable between low- and high-confluence conditions (Fig. S6E). In addition, earlier work has demonstrated that anoxic or hypoxic stress can induce hepatocyte vacuolation (Sykes et al., 1976). Consistent with this, hypoxic conditions in the present study triggered pronounced vacuolation accompanied by increased Rab38 expression (data not shown). Taken together, these findings suggest that LRO remodeling in hepatocytes is responsive to specific forms of cellular stress. Such remodeling may be particularly relevant under pathological conditions, including hepatitis, acute metabolic dysfunction, metabolic-associated steatotic liver disease (MASLD) or hepatocarcinogenesis; conversely, alterations in LRO organization may also contribute to the development or progression of these pathological states. Further mechanistic studies will be required to define the upstream regulators and to establish the physiological and pathological relevance of stress-induced LRO remodeling in hepatocytes.

Although the redundant roles of Rab32 and Rab38 are well recognized, this study provides evidence suggesting that these two Rab proteins may also exert differential functions. Rab32 and Rab38 contributed to LD homeostasis in distinct, yet complementary, ways: Rab32 depletion resulted in enlarged LDs, whereas loss of Rab38 preferentially led to the accumulation of numerous small LDs (Fig. S3). This divergence in LD morphology closely resembles phenotypes associated with impaired cytosolic lipolysis versus defective lysosomal LD degradation, respectively (Schott et al., 2019).

These observations raise the possibility that Rab32/38-positive LROs act at different stages of LD turnover. Rab32 may primarily influence early LD remodeling steps that limit LD expansion, whereas Rab38 may function at a downstream stage required for efficient clearance of smaller LDs. Consistent with these functional differences, LD incorporation into acidified or LAMP1-positive compartments was markedly reduced in *Rab38*-deficient and *Rab32/38* double-knockdown cells compared with *Rab32*-deficient cells.

This study has several limitations that also highlight important avenues for future work. The *in vivo* analyses were performed using whole-body *Rab32/38* double-knockout mice rather than hepatocyte-specific models, meaning systemic influences cannot be excluded; generating liver-specific conditional knockouts and examining primary hepatocytes will be essential to establish the physiological relevance of the Rab38-VPS4B axis. Imaging limitations also constrained our ability to visualize early LD invagination events, which appeared far more transient than membrane fusion; future high-speed or super-resolution approaches such as lattice light-sheet or correlative light-electron microscopy may help resolve these early steps with greater precision. Furthermore, while reduced LD intensity in *Vps4a/b*-deficient cells suggests compensatory activation of ATGL-driven lipolysis, direct biochemical measurements of ATGL expression and activity will be required to clarify its hierarchical relationship with the LRO pathway. Finally, the hypoxia-induced vacuolation and Rab38 upregulation observed here warrant deeper characterization. Future studies should evaluate whether glycogen content, stress-responsive transcription factors (e.g., HIF family), or pathological stimuli modulate this response, and whether similar regulatory principles operate in primary hepatocytes or *in vivo* under metabolic stress.

## Acknowledgments

This study was funded by Grants-in-Aid for Scientific Research KAKENHI (20H05326, 22H04647, 23H02475) to T.N. and (20K15789, 25K09629) to S-L.L., Hirose Foundation to S-L.L. and (JPMJSP2138) to Z.Z.

## Author contributions

T.No. and S-L.L. supervised this study. Z.Z. performed the experiments and wrote the first draft. S-L.L., Y.K., B.C., T.Z., Y.L., Y.U. assisted the experiments. T. Ni, R.S. T.K, N.U, S.T. provided resources. All authors reviewed the results and approved the final version of the manuscript.

---

**Supplementary Figure 1. Rab32/38-positive LROs are distinct from conventional lysosomes in AML12 cells.**

**A.** Representative fluorescence images of AML12 cells expressing GFP-Rab32 or GFP-Rab38 together with LAMP1-RFP under standard culture conditions. Arrows indicate LAMP1-RFP positive and GRP-Rab32/38 negative signaling. Scale bar, 5 µm. **B.** Pearson’s correlation coefficients measuring colocalization between GFP-Rab32 or GFP-Rab38 and LAMP1-RFP under vehicle and orlistat conditions (50 µM, 24 h). Data are mean ± SD; each dot represents one randomly selected field (n = 5 per condition). **C.** Quantification of vacuole number (>3 µm²) per field in WT, sh*Rab32*, sh*Rab38*, and DKD AML12 cells (n = 10 random fields per condition, 5 cells per field). Data are presented as mean ± SD. Statistical significance was determined using the Kruskal-Wallis test with multiple comparisons (WT vs sh*Rab32*, ns; WT vs sh*Rab38*, ns; WT vs DKD, *p* = 0.0117). **D.** Representative fixed-cell fluorescence images of AML12 cells expressing GFP-Rab32 and stained with DAPI (blue) and Lipi Deep-Red (red) under vehicle or oleic acid loading (200 µM, 24 h). Arrowheads indicate lipid droplets internalized within invaginating structures of Rab32-positive LROs. Scale bar, 2.5 µm.

**Supplementary Figure 2. Differential association of lipid droplets with LC3-positive autophagic structures and Rab32/38-positive LROs in AML12 cells.**

**A.** Immunoblot analysis of autophagy-related markers in AML12 cells cultured under different nutrient and confluence conditions. AML12 cells were maintained at low or high confluence and treated with or without Bafilomycin A1 (Baf A1; 200 nM, 2 h), as indicated. Protein levels of p62 and LC3-II were analyzed by immunoblotting. Tubulin was used as a loading control. **B.** Representative fluorescence images of AML12 cells under steady-state culture conditions, showing the spatial relationship among lipid droplets (Lipi-Blue), LROs (GFP-Rab38), and LC3-positive structures (mRFP-LC3). Scale bar, 2 µm. **C.** Schematic illustration of macroautophagy and its inhibition by excess Atg4B^C74A^. **D.** Validation of the macroautophagy-blocking function of mStrawberry-Atg4B^C74A^. AML12 cells were transiently transfected with the mutant and subjected to nutrient starvation, followed by LC3 immunostaining. Dashed lines outline two adjacent cells: Cell 1, expressing mStrawberry-Atg4B^C74A^, shows markedly reduced endogenous LC3 puncta; Cell 2, a non-transfected neighbor, displays robust LC3 puncta formation. Scale bar, 5 µm.

**Supplementary Figure 3. Rab32/38 deficiency alters lipid droplet size distribution in AML12 cells.**

**A-D.** Analysis of lipid droplet (LD) size distribution in control, sh*Rab32*, sh*Rab38*, and *Rab32/38* double-knockdown (DKD) AML12 cells. **A.** Violin plots showing the distribution of average LD size, with red lines indicating the median. **B-D.** Binned LD frequency distributions, presented as the relative percentage of LDs within each size category.

**Supplementary Figure 4. Rab32/38 double knockdown impaired LRO-mediated LD engulfment but did not affect autophagic flux under basal conditions.**

**A.** Schematic illustration of the analysis of microlipophagy using a tandem-fluorescent perilipin reporter. N-terminal fusion of mCherry (mChe) and GFP to PLIN2 labels the surface of cytoplasmic LDs. Upon delivery of LDs into acidic organelles, such as lysosomes, the GFP signal is selectively quenched, resulting in puncta exhibiting mCherry fluorescence only. The degree of colocalization between mCherry and GFP reflects the level of cytoplasmic LDs. **B.** Representative fluorescence images of scramble and *Rab32/38* DKD AML12 cells expressing the mCherry-GFP-PLIN2 dual-color LD reporter, used to assess LRO-mediated LD engulfment efficiency. Scale bar, 5 µm. **C.** Quantification of mCherry-GFP colocalization using Pearson’s correlation coefficient in scramble versus DKD cells (n = 23 cells; Welch’s t-test, *p* = 0.0337). Data are presented as mean ± SD. **D.** Immunoblotting and in-gel fluorescence detection of total cell lysates from scramble and *Rab32/Rab38* DKD AML12 cells stably expressing Halo-LC3B. Cells were pulse-labeled for 20 min with 100 nM tetramethylrhodamine (TMR)-conjugated ligand in nutrient-rich medium, immediately harvested (0 h), or incubated in starvation medium for 6 h. Where indicated, cells were incubated in starvation medium in the presence or absence of 100 nM bafilomycin A1 (BafA1) for the final 2 h before collection.

**Supplementary Figure 5. PI(3,5)P₂ contributes to LRO membrane dynamics required for LD uptake.**

**A.** Live-cell fluorescence images of AML12 cells expressing the PI(3,5)P₂ reporter PX-SnxA^GV^-GCC-mCherry together with GFP-Rab38. Cells were imaged with or without 100 nM apilimod (AP) treatment for 30 min. Scale bar, 5 µm. **B.** Live-cell imaging after AP washout, performed using the same experimental scheme as in **A**, followed by orlistat treatment to prevent LD degradation. Images were acquired at 24 h after AP washout to visualize PX-SnxA^GV^-GCC-mCherry and GFP-Rab38 signals together with LDs under 24-h orlistat treatment. Scale bar, 5 µm.

**Supplementary Figure 6. Supporting metabolic and molecular characterization of Rab32/38-deficient models.**

**A.** Berlin blue staining of livers from 1-year-old WT and *Rab32/38* DKO mice. Berlin blue staining was performed to assess iron deposition in liver sections (WT, n = 3; DKO, n = 3). Scale bars: 25 µm. **B-C.** Serum and liver triacylglycerol (TAG) levels in HFD-fed male mice. **B**. Serum TAG levels. **C.** Liver TAG levels. Hepatic lipids were extracted using the Folch method and quantified using a commercial enzymatic TAG assay kit. Each data point represents an individual mouse. Data are presented as mean ± SD (WT, n = 3; DKO, n = 3). Statistical significance was determined using an unpaired Student’s t-test; ns indicates p > 0.05. For liver TAG levels, *p* = 0.0570. **D.** Immunoblot analysis of Rab32 and Rab38 expression in AML12 cells and liver tissues from WT and DKO mice. Tubulin was used as a loading control. Asterisks indicate the specific bands corresponding to Rab32 and Rab38 at the expected molecular weights. **E.** Immunoblot analysis of phosphorylated S6 kinase (p-S6K) and total S6K levels in AML12 cells cultured under low- and high-confluence conditions. Tubulin was used as a loading control. Band intensities of p-S6K and total S6K were each normalized to tubulin, and the p-S6K/S6K ratio was calculated based on the normalized values. The quantified p-S6K/S6K ratios are indicated below the blots. Molecular weight markers are shown on the right.

