## Supplementary figures and images for "Rab32/Rab38-positive Lysosome-Related Organelle degrades lipid droplet in hepatocytes by microautophagy"

### Supplementary Figures

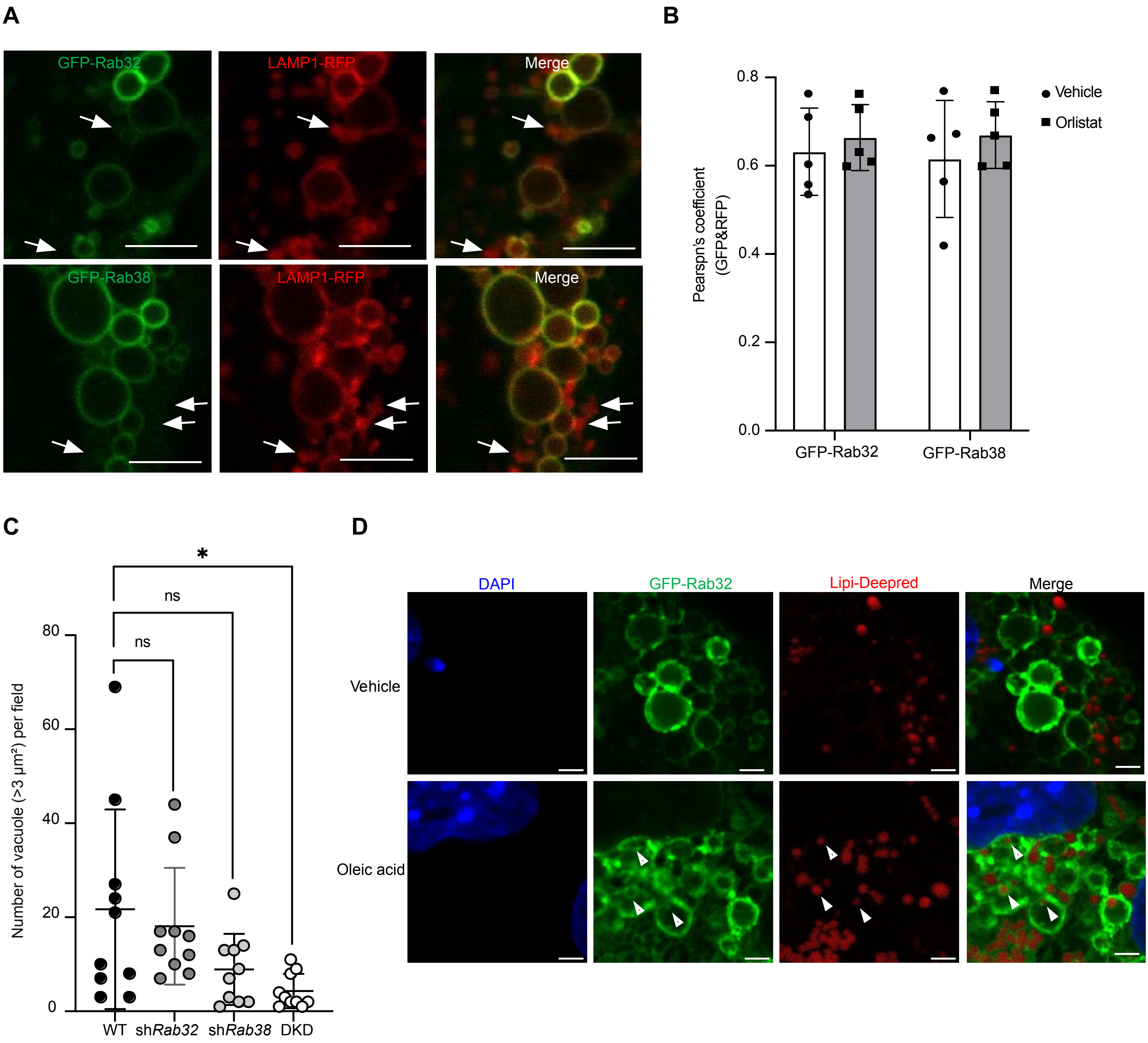

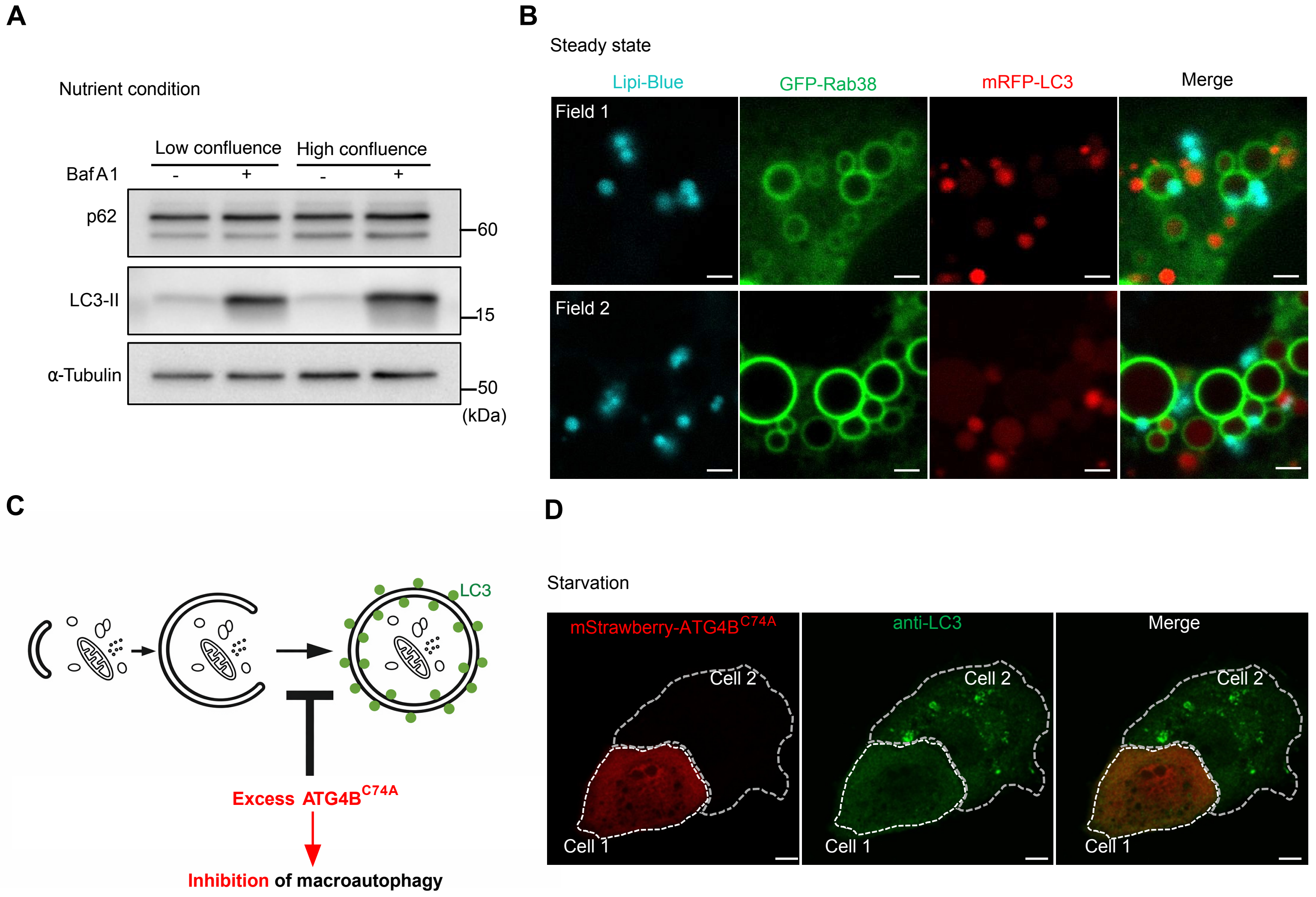

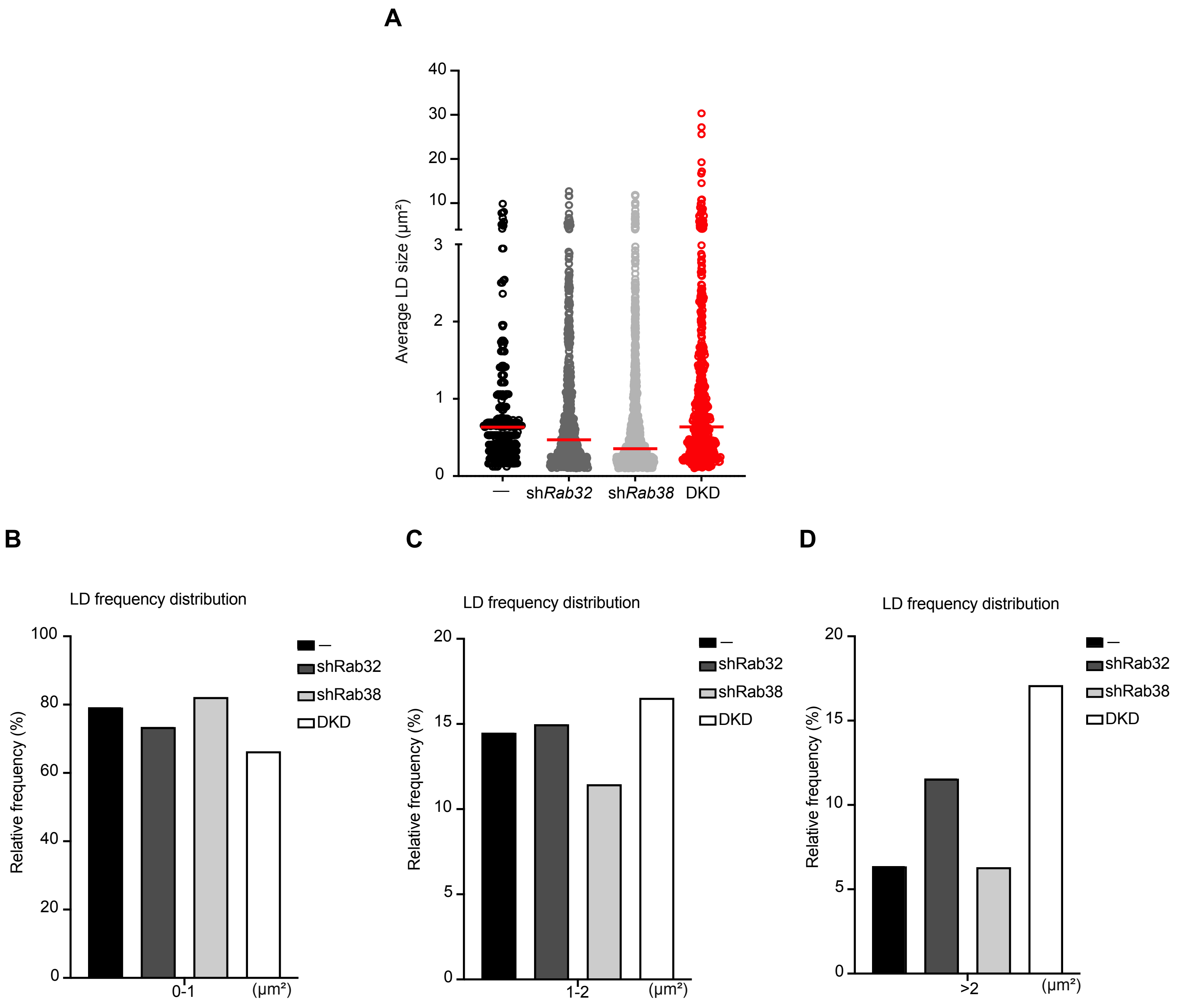

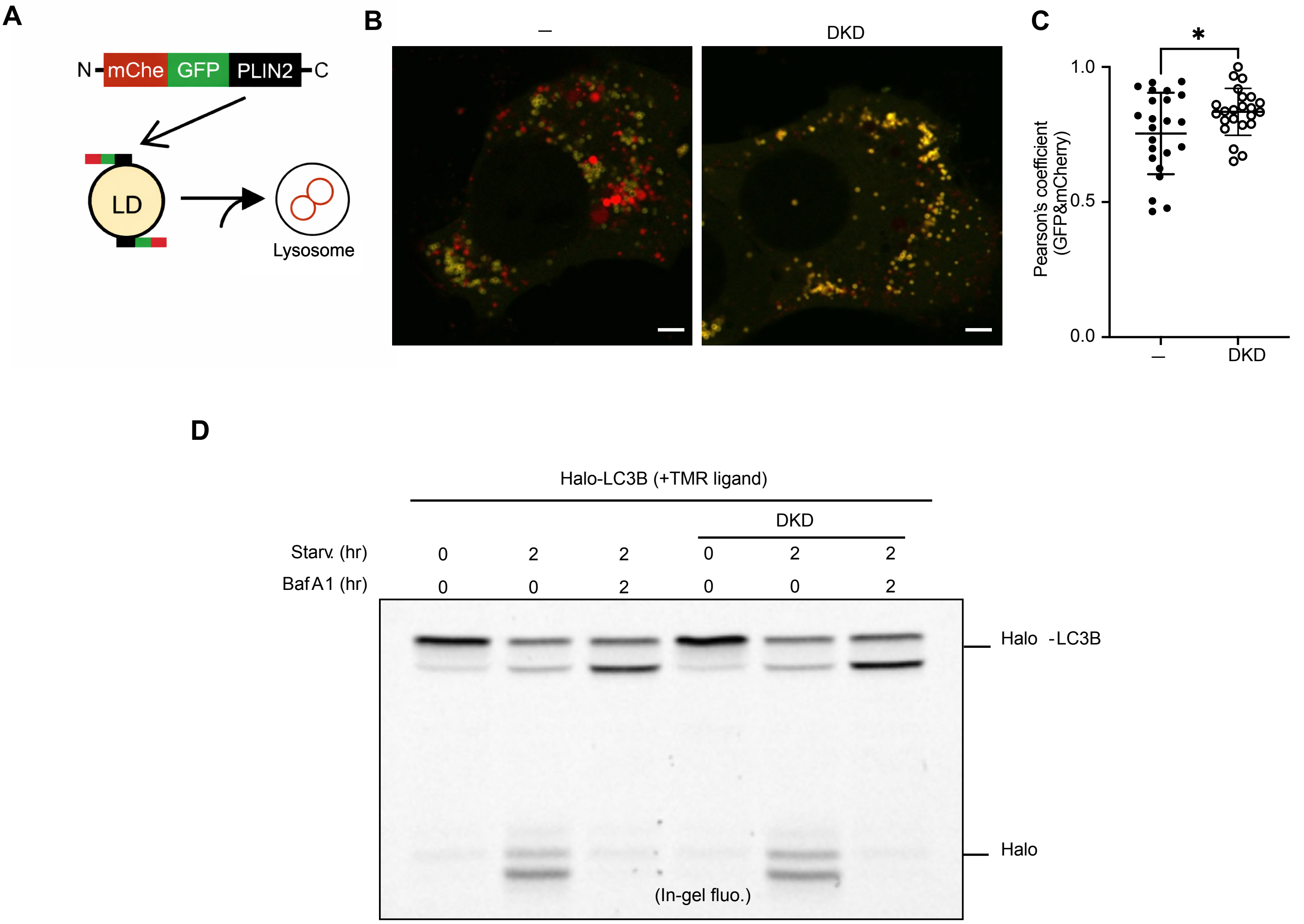

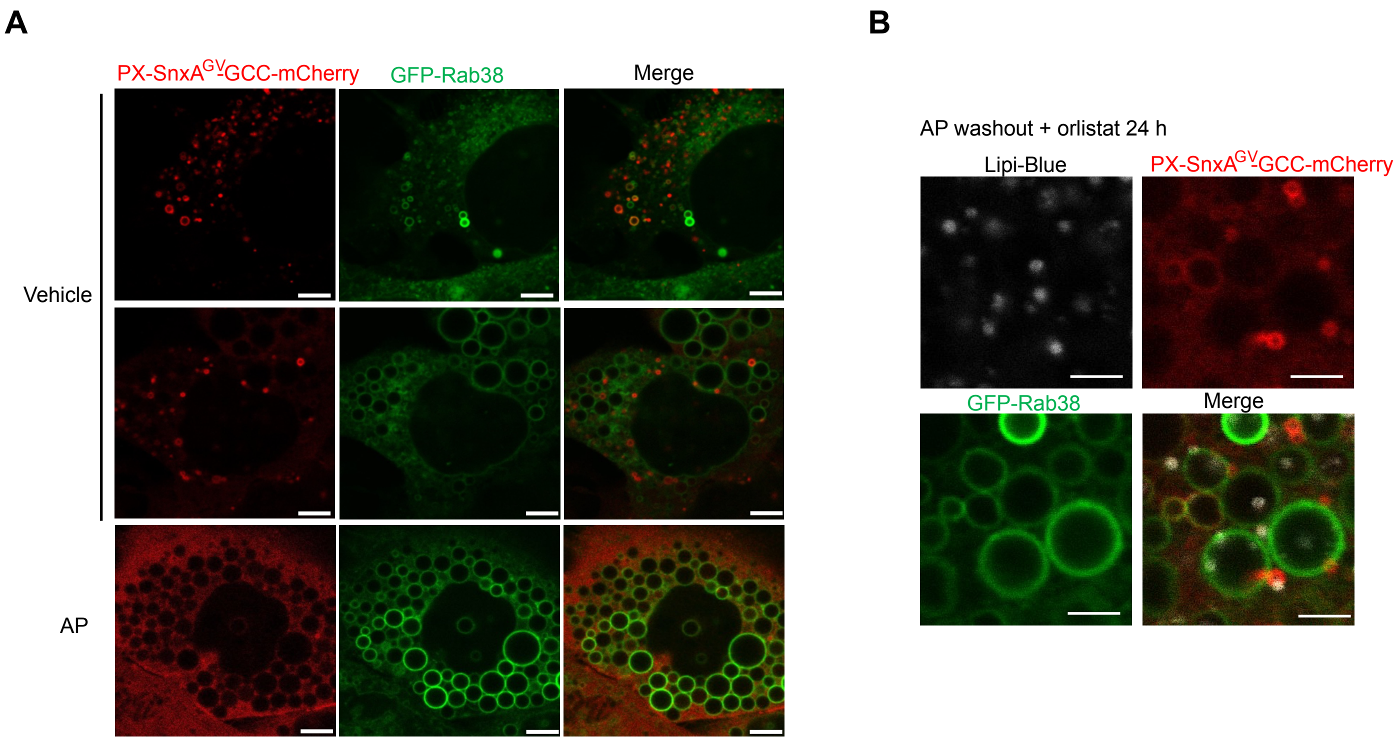

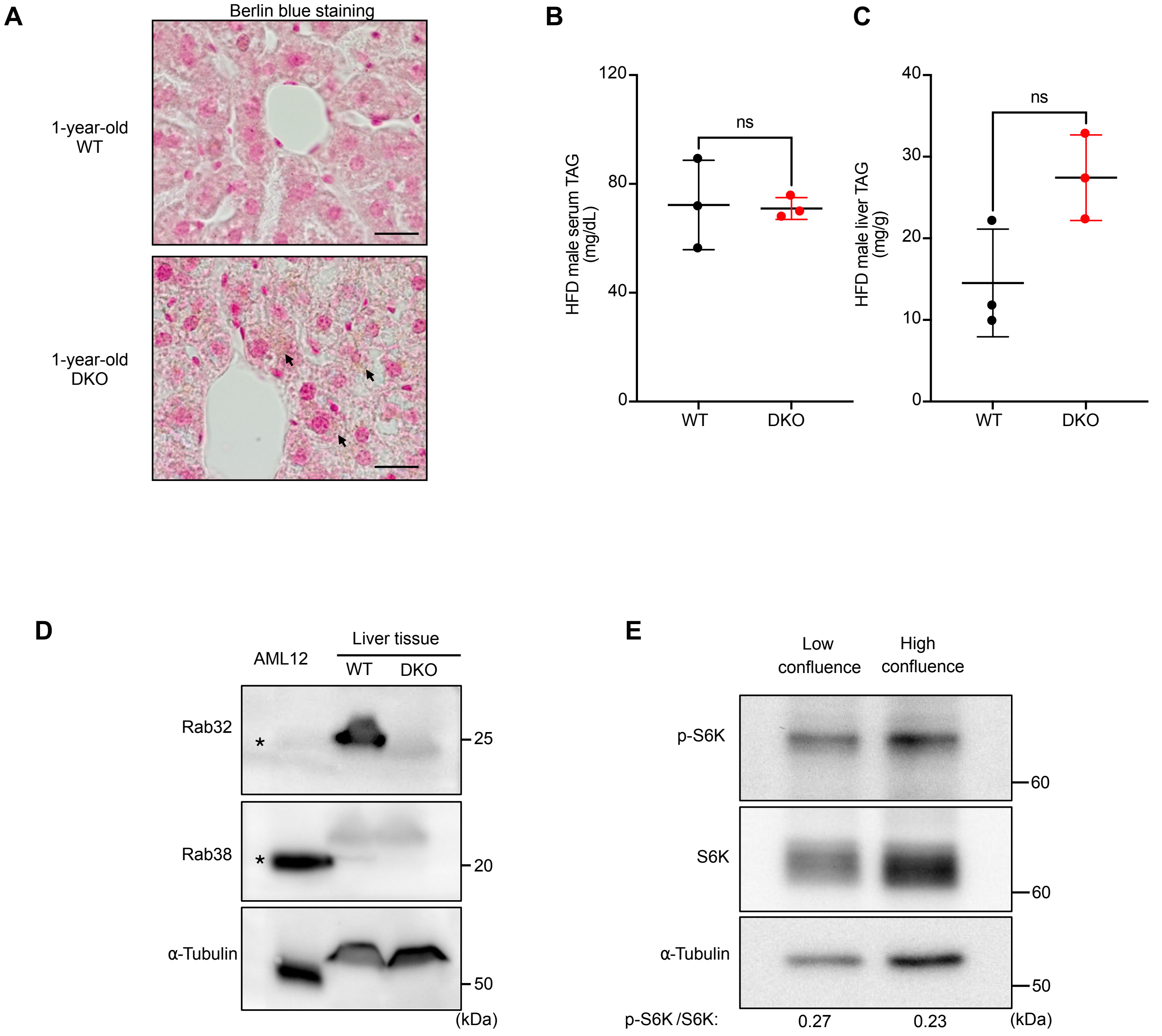
